# Shark-dust: High-throughput DNA sequencing of processing residues unveils widespread trade in threatened sharks and rays

**DOI:** 10.1101/2022.12.16.520728

**Authors:** Andhika P. Prasetyo, Joanna M. Murray, Muh. Firdaus A. K. Kurniawan, Naiara G. Sales, Allan D. McDevitt, Stefano Mariani

**Author notes:** Corresponding author. A. P. Prasetyo (,) and S. Mariani. These authors contributed equally (Allan D. McDevitt, Stefano Mariani).

## Abstract

Illegal fishing, unregulated bycatch, and market demand for certain products (e.g. fins) are largely responsible for the rapid global decline of shark and ray populations. Controlling trade of endangered species remains difficult due to product variety, taxonomic ambiguity and trade complexity. The genetic tools traditionally used to identify traded species typically target individual tissue samples, are time-consuming and/or species-specific. Here, we performed high-throughput sequencing of trace DNA fragments retrieved from dust and scraps left behind by trade activities. We metabarcoded ‘shark-dust’ samples from seven processing plants in the world’s biggest shark landing site (Java, Indonesia), and identified 54 shark and ray taxa (representing half of all chondrichthyan orders), half of which could not be recovered from tissue samples collected in parallel from the same sites. Importantly, over 80% of shark-dust sequences were found to belong to CITES-listed species. We argue that this approach is likely to become a powerful and cost-effective monitoring tool wherever wildlife is traded.

**One-Sentence Summary:** Shark-dust, the traces of biological material left behind from the processing of shark products, can now be DNA-sequenced in bulk to accurately reconstruct the biodiversity underlying trade.

## Introduction

Continued and increasing anthropogenic stressors have devastated habitats and wildlife across the globe, including the dramatic depletion of sharks and rays (hereafter referred to as ‘elasmobranchs’) (Dulvy et al. 2021). Conservative life-histories (Mardhiah et al. 2019) make elasmobranchs vulnerable to fisheries overexploitation, and their extirpation can destabilise functional diversity and ecosystem structure (Dulvy et al. 2021). Although some elasmobranch fisheries can be sustainably managed (Simpfendorfer and Dulvy 2017), the market demand for high value products, such as fins, liver oil and gill plates, typically leads to overexploitation of elasmobranch resources (Dulvy et al. 2021), which is then further fuelled by illegal and unreported catches.

This combination of market demand, over-exploitation, and lack of detail in catch and trade data (Cawthorn et al. 2018) requires effective mechanisms to monitor elasmobranch populations and ensure their sustainable management (Prasetyo et al. 2021). This includes improved catch reports, special regulations for endangered species (e.g. the Convention on International Trade in Endangered Species of Wild Fauna and Flora (CITES, (Pavitt et al. 2021)), and a range of other transdisciplinary initiatives (Booth et al. 2019). A fundamental critical step in this context is the accurate reconstruction of the biodiversity composition of elasmobranch products at landing sites, processing plants, markets and export hubs.

This year, the difficulty of the task has more than tripled, as the number of CITES-listed species has increased from 47 to 151 (CITES 2022a); yet, species listed in Appendix II can still be traded, by taking into account the viability of exploitation within the context of the Non-detrimental Findings (NDF) framework (Smith et al. 2011). Thus, conservation managers now face a scenario where 14% of the 1,120 described elasmobranch species (nearly one third of which deemed to be under some level of conservation threat, (IUCN 2021) can still be traded and substituted for other species under greater restrictions. Understanding and regulating trade in these species is challenging because elasmobranch products are extremely diverse in both their usage and their value, and are processed in a myriad of different ways (Dent and Clarke 2015). Due to their similarity in appearance and lack of distinctive features in most derivative products, shark and ray species can be deliberately or accidentally mislabelled by those involved in the trade (**Figure 1**). This has led to the rapid development of molecular technologies, which progressively made DNA-based inference a staple of wildlife forensics. Of these, DNA barcoding (Shivji et al. 2002) and mini-barcoding (Fields et al. 2015) can robustly identify species in fresh and processed samples, while real-time qPCR (Cardeñosa et al. 2018), LAMP-based (But et al. 2020) and universal close-tube barcoding (Prasetyo et al. 2022) assays can detect target species in a matter of hours.

**Figure 1.**
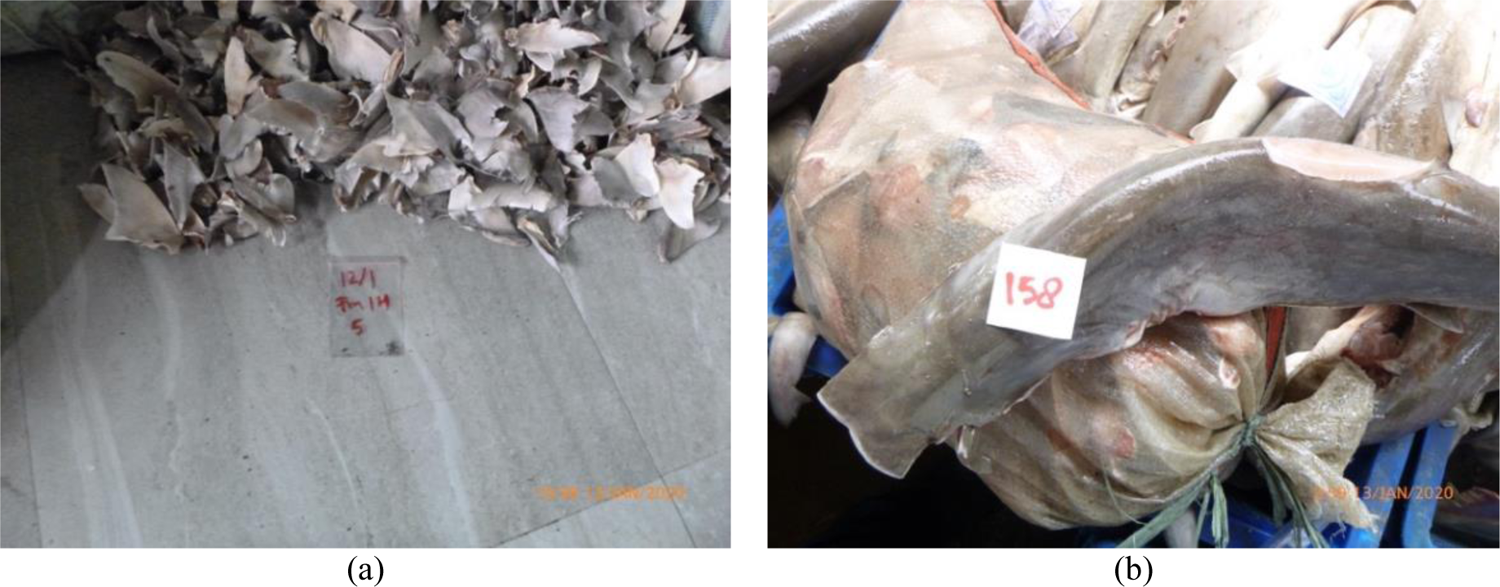
Condition of sample collection for (a) shark-dust from a pile of small dried fins, and (b) tissue sample from a finless juvenile scalloped hammerhead shark whose cephalofoil (the distinctive “face” in this Family, also known as “blade”) had been cut.

However, all these methods require the collection and analysis of individual specimens, which is a significant limitation when large volumes of samples, across many locations, must be inspected in a limited timeframe to estimate species composition and detect species under trading restrictions (Prasetyo et al. 2021). Recent advances in next generation sequencing (NGS) have shaped the transformation of general DNA barcoding (Hebert et al. 2003) into a technique that allows the simultaneous identification of multiple taxa from an inordinate mixture, known as DNA metabarcoding (hereafter referred to as just ‘metabarcoding’) (Riaz et al. 2011). These principles have been broadly applied to analysing environmental DNA (eDNA) samples; trace DNA fragments left behind by organisms in water, soil and air, an approach that effectively complements – and in some cases surpasses – traditional monitoring, especially when labour and expertise are scarce (Boussarie et al. 2018). Such developments are unlocking novel applications in trade monitoring, allowing bulk mixtures to be analysed and tackling the limitations of existing tools.

Here we propose a novel metabarcoding application, by targeting seven key shark and ray trading hubs in the island of Java, Indonesia, the top elasmobranch-landing country in the world. We used high-throughput metabarcoding to screen the by-products of processing plant activities (which we term ‘shark-dust’) and compare them with single-specimen barcoding. This unconventional application is poised to minimize labour requirements, enhance the detection of species that are not visible at the time of inspection, and be implemented globally.

## Materials and methods

### Study sites

Indonesia’s geographical location and its vast and complex coasts make it a unique and emblematic marine megadiversity hotspot. Between 2007 and 2017, Indonesia was the top elasmobranch landing country (Okes and Sant 2019) but export statistics revealed substantial knowledge gaps and inaccuracies (Prasetyo et al. 2021). Here we targeted seven locations across cities on Java Island, the most populous island in Indonesia (**Figure S1**) and the main export hub for various export commodities, including elasmobranch products. The locations included elasmobranch processing plants (PP), export hubs (EH) and an inspector station (AU).

### Sample collection

Dust and tissue samples were collected from January to February 2020. We collected two sets of samples: first, we gathered 28 mixtures of residual material from floors and surfaces where shark products were processed, sorted, and stored for later shipping, henceforth referred to as “dust” samples (**Table S5**); then, we selected 183 tissue samples from individual specimens (**Table S6**). Replicated samples (4 ±3 samples) were collected in seven locations representative of Indonesia’s processing, export, and regulatory activity. About 10 grams of dust were scooped and stored at room temperature in sterilised 5 ml Click-Seal flat bottom tubes without a preservative. From the same location, about 10 g of tissue was collected from individual specimens opportunistically found at the sites, including both fresh and processed products. The tissue was then stored in 2.0 mL screw-cap microcentrifuge tubes, submerged in 90% ethanol and stored at 4°C.

### Laboratory procedures

DNA was extracted from all samples (dust and tissue samples) following the Mu-DNA protocol for tissue samples (Sellers et al. 2018) with an overnight incubation and a final elution volume of 100 μl. Dust samples were stored in the sealed bag at room temperature and were handled using sterile instruments. We also processed 183 tissue samples from the same locations where dust samples were collected. Tissue samples were extracted similarly to the dust samples. All DNA extractions were diluted to 10-15 ng/μl prior to DNA amplification. The Elas02 primer pairs (Elas02-F, 5’-GTTGGTHAATCGTGCCAGC-3’; ElasO2-R, 5’-CATAGTAGGGTATCTAATCCTA-GTTTG-3’) was used to target a ~180 bp amplicon from a variable region of the 12S rRNA mitochondrial gene (Miya et al. 2015; Taberlet et al. 2018). Given that dust was sampled from the floor, an elasmobranch-specific 12S marker was selected to avoid non-target amplification, as the use of a COI-based marker would likely lead to the vast majority of reads coming from other organisms (Collins et al. 2019). Samples were amplified in triplicate to minimize amplification stochasticity, and replicates were later pooled into a single representative sample. Meanwhile, the sequencing of individual tissue samples followed a massively parallel framework hereafter termed ‘high-throughput barcoding’ (HTB). The Leray-XT primer pair targeting a ~313 bp amplicon from a region of the COI mitochondrial gene (Wangensteen et al. 2018) was used for DNA amplification from tissue samples.

Adapters were ligated to PCR products using the KAPA Hyper Prep Kit PCR-Free protocol with incubation time at 7 minutes and bead clean at a 0.9 ratio. Libraries were then quantified by qPCR using the NEBNext^®^ Library Quant Kit for Illumina sequencing. The dust-generated library was diluted to 6 nM, and sequenced on an Illumina MiSeq run using a 2×150 bp v2 kit; the tissue sample libraries were diluted to 4 nM and sequenced in one Illumina MiSeq run using a 2×300 bp v3 kit. PhiX spike was at 1% for both runs.

### Bioinformatics and statistical analysis

Bioinformatic analysis was carried out using the OBITools metabarcoding package (Boyer et al. 2016) and the taxonomic assignment was conducted using ecotag against a custom reference database (**Figures S2-S3, Table S7**). To obtain an accurate estimate of occurrence (Deagle et al. 2019) and correct for both the exponential nature of PCR in the dust samples and the unknown bulk of the different species along the processing stages, a square root transformation and relative read abundance (RRA) metric were applied. Sampling effort and sample types were evaluated with species accumulation curves plotted with the R package BiodiversityR (Kindt and Coe 2005) using the ‘exact’ method. To evaluate compositional differences between sampling techniques, we converted species detections from both data sets into presence-absence data by locations, then calculated one dissimilarity index (Jaccard, for binary MOTU data) whose configuration was visualised via multidimensional scaling, using the function ‘metaMDS’. We also formally tested differences between shark-dust and tissue samples with a PERMANOVA (999 permutations) using the function ‘adonis’. Both functions were run in the R package VEGAN (Oksanen et al. 2013). Statistical analyses were performed in the R program environment (R Development Core Team 2012, version 3.6.0). Further details on laboratory and bioinformatic procedures can be found in the **Supplementary Materials**, and the scripts associated with the study are provided at: https://github.com/andhikaprima/sharkdust.

## Results and discussion

### Dust metabarcoding analysis

We obtained around 5.6 million reads from 28 discrete dust samples. We refined the final dataset to 4,640,239 elasmobranch-only reads, partitioned into 61 MOTUs (**Figures S1-S2, Figure S5, Table S1**) belonging to seven different orders: Carcharhiniformes, Lamniformes, Squaliformes, Hexanchiformes, Orectolobiformes, Myliobatiformes, and Rhinopristiformes. Taxonomic assignment successfully identified 54 of the 61 MOTUs to species level, with five assigned to genus level and two only attributable to families.

Nearly 84% of the total reads belonged to 32 CITES-listed taxa, including high profile pelagic bycatch species, such as hammerhead sharks (*Sphyrna* spp.), silky shark (*Carcharhinus falciformis*) and spot-tail shark (*Carcharhinus sorrah*) (**Figure 2a**). The scalloped hammerhead shark (*S. lewinĩ*) could be found almost everywhere, but it was most prevalent in the processing plants in Indramayu (IDM2 and IMD3), Banyuwangi (BYW7), and Surabaya (SBY6). The spottail shark, recently added to the CITES list, had a substantial presence (in terms of read abundance) in dust samples collected in the Indramayu processing plants (**Figure 2b**). Among non-CITES-listed species, tiger shark (*Galeocerdo cuvier*) was the predominant species across sampling locations, followed by zebra shark (*Stegostoma fasciatum*), the Australian weasel shark (*Hemigaleus australiensis*), whitespotted whipray (*Himantura gerrardi*) and spotless smooth-hound (*Mustelus griseus*) (**Figure 2c**). These five species contributed about 70% of the non-CITES-listed read count overall, but their relative proportions varied greatly among locations.

**Figure 2.**
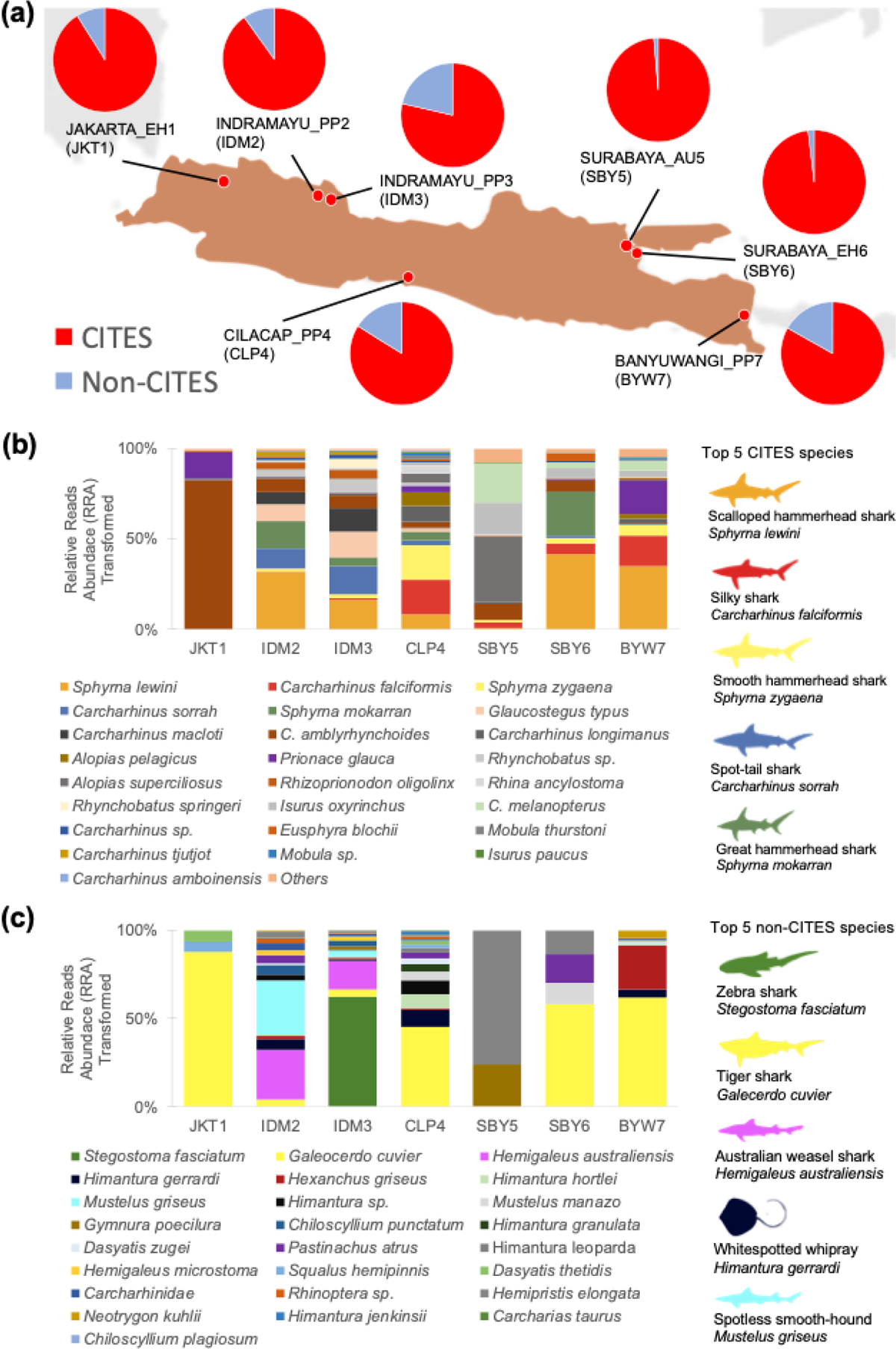
CITES and non-CITES listed species composition (in square-rooted read abundance) across sampled locations (a); composition of CITES-listed species (b), and composition of non-CITES-listed species (c). Top-5 species are visualized with silhouettes and same colour in the bar chart after normalized. Read abundance values were square-root transformed.

The prevalence and abundance of reads from CITES-listed species detected in dust samples shows that these animals continue to be major trade commodities and that monitoring efforts need to be intensified. Such species of conservation concern – primarily pelagic taxa – are found in abundance in processing plants (IDM2, IDM3, CLP4 and BYW7) and exporter warehouses in main export hub cities (i.e. Jakarta and Surabaya (JKT1 and SBY6)). These results amplify earlier indications that CITES-listed species, such as thresher sharks, hammerhead sharks, silky shark, wedgefishes, and guitarfishes, are still being traded in major Indonesian markets (Fahmi et al. 2021) and may still be exported through Non-Detrimental Finding (NDF) mechanisms (CITES 2022b). In Hong Kong, which is the main destination market, fin products of CITES-listed species are still frequently traded (Okes and Sant 2019) and modelled to be ~10% of the overall traded volume (Fields et al. 2017). Based on our results from the world’s largest exporter – and the recent expansion of CITES listings – these figures are likely an underestimation. Dust samples also detected several key reef-associated sharks as trade commodities, such as blacktip reef shark (*C. melanopterus*), whitetip reef shark (*Triaenodon obesus*) and sand tiger shark (*Carcharias taurus*). These species play an important part in the equilibria of coral reef ecosystems, which is particularly concerning for Indonesia, where reef-sharks have been driven to near functional extinction (MacNeil et al. 2020). Several mesopredators among the rays were also detected, including Hortle’s whipray (*Himantura hortlei*), mangrove whipray (*Himantura granulata*), pale-edged stingray (*Dasyatis zugeĩ*), and bluespotted stingray (*Neotrygon kuhlii*). These species, albeit not controlled under CITES, significantly contribute to trophic interactions in key coastal ecosystems (Flowers et al. 2021); in fact, 90% of non-CITES-listed species detected from dust samples are currently designated as threatened species (Near Threatened, Vulnerable, Endangered or Critically Endangered) under the IUCN (International Union for Conservation of Nature) Red List (IUCN 2021). Therefore, beyond trade enforcement aspects, obtaining information on these taxa is critical for monitoring the impact of exploitation on population dynamics and ecosystem health.

### Comparison of species detections from dust and tissue samples

Tissue-based barcoding successfully identified 175 out of 183 samples associated with the locations where dust samples were taken. Specimens were partitioned into 36 taxa, nearly all of which were also detected in the dust samples (**Figure 3a**). Overall, we were able to identify more than 70 taxa across methods; however, the dust samples detected 16 more genera than tissue samples and identified 11 unique CITES-listed species **(Figure 3b, Figure S4, Table S2**). When sequencing reads from the dust samples were transformed into presence and absence data, species compositions between dust and tissue samples were shown to be significantly different (PERMANOVA: F=3.49, p=0.001; **Figure 3c, Table S3**). Tissue samples show a greater separation among locations, due to the high-grading bias introduced by the single-specimen approach to sampling (which may also select for more ‘notable’ samples). Dust samples showed a consistently greater alpha diversity across locations, detecting an average of 31.57 (±16.34) taxa per sample, with tissue samples averaging 11.14 (±6.01), as is also shown by the taxon accumulation curve (**Figure 4a**).

**Figure 3.**
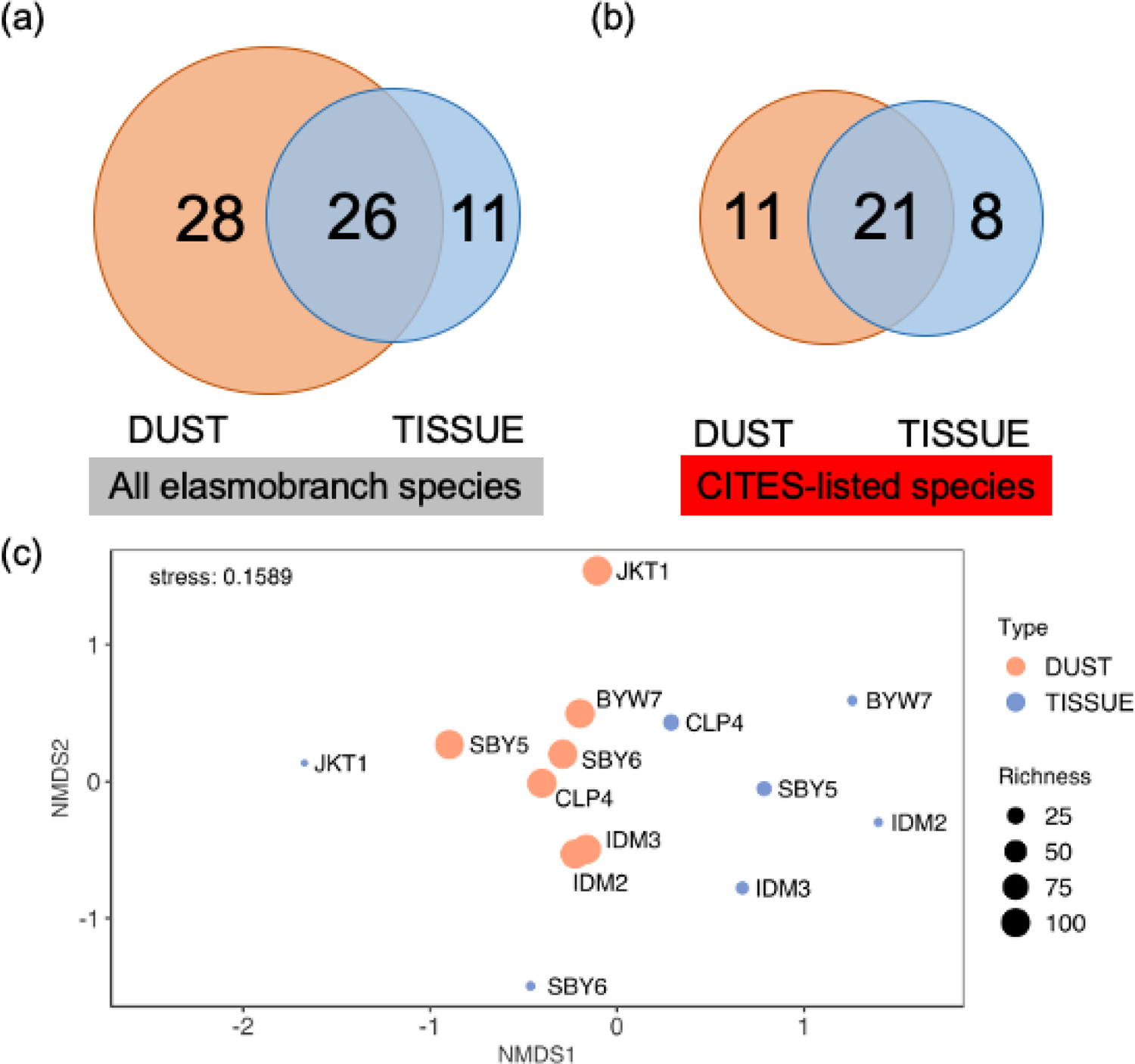
Comparison between species recovery from dust and tissue samples; Venn diagrams of all elasmobranch species (a), CITES-listed species only (b), and non-metric multidimensional scaling (nMDS) based on Jaccard similarity index between two sample types in different locations (c). Samples have been pooled into the 7 locations. Nb. Only species-level taxa are considered except for Mobula sp. and Rhynchobatus sp. as these taxa were detected by dust metabarcoding, despite the 12S marker being unable to discriminate between closely related species in these genera.

**Figure 4.**
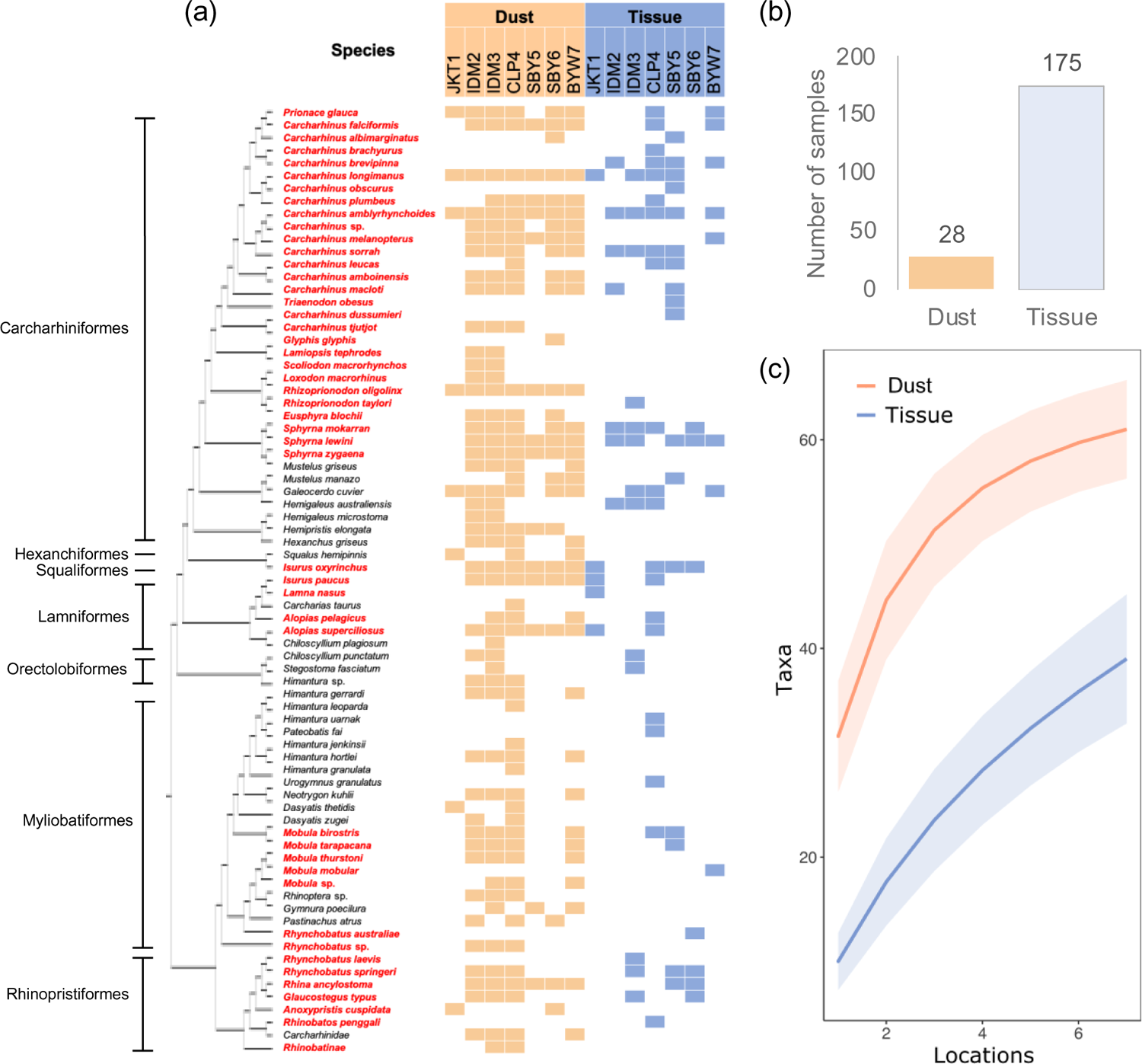
The cladogram (a) was generated using FigTree 1.4.4 using NADH2 region sequences (Naylor et al. 2012) from the NCBI database. Colours represent sample type, such as dust samples (ORANGE) and tissue samples (BLUE) for results from each sampling location, with CITES-listed species written in RED. Species accumulation curves (b) emphasize the differences in alpha diversity recovery between methods.

Dust metabarcoding has much greater power to unveil a comprehensive portrayal of shark and ray species being traded, for a considerably lower sampling effort (N_dust_= 28 *vs* N_tissue_= 175) and less disruption of the processing and trading operations in the visited hubs (**Figure 4b**). Dust samples revealed some cryptic and rare species, such as winghead shark (*Eusphyra blochiĩ*),pigeye shark (*C. amboinensis*), sand tiger shark (*Carcharias taurus*), smooth hammerhead (*S. zygaena*), knifetooth sawfish (*Anoxypristis cuspidata*), manta and devil rays (*Mobula* spp.). The latter three are hardly ever seen at landing places, given their fully protected status under Indonesia’s regulations. These findings mirror the performance of eDNA studies on elasmobranchs from natural environments, which consistently reveal important ‘dark diversity’ that is missed by pre-existing biomonitoring tools (Boussarie et al. 2018). In this sense, the ‘shark-dust’ metabarcoding approach can boost and streamline all the biodiversity, fishery, and trade control operations that have up to this point been carried out via earlier-generation DNA monitoring tools.

There were 40 CITES-listed taxa identified in total, with 12 taxa, including thresher sharks (*Alopias* spp.), mako sharks (*Isurus* spp.) and two hammerhead species that are commonly found at landing sites (*S. lewini* and *S. mokkaran*) identified using both dust and tissue samples. Meanwhile, tissue samples revealed one species that is not distributed in Indonesian waters, i.e. porbeagle shark (*Lamna nasus);* but this was a sample obtained from the exporter’s reference collection that was used for education purposes.

### A cutting-edge tool for trade monitoring

Our findings showed that trade monitoring using dust metabarcoding expands the reach of traditional barcoding methods. However, seven (7) MOTUs could not be identified to species level from dust samples (**Table S4**), including two families and five genera with species listed in CITES appendices, namely wedgefishes (*Rhynchobatus* sp.), devil rays (*Mobula* sp.) and requiem sharks (*Carcharhinus* sp.) and guitarfishes (Rhinobatinae). We had anticipated this issue by developing an additional 12S reference database for our analyses, but recent studies (Mariani et al. 2021; Miya et al. 2020) had already shown that the size (170-180bp) and resolution of the 12S Elas02 fragment will not allow discrimination between some closely related species, as shown for *Rhynchobatus, Mobula*, Rhinobatinae, and also for some species in the taxonomically problematic and polyphyletic genus *Carcharhinus* (Sorenson et al. 2014). Yet, despite these limitations, the marker used remains the most effective metabarcoding tool for elasmobranch identification whilst also avoiding non-target amplification (Collins et al. 2019), and this could be further strengthened through the ongoing expansion of 12S and mitogenomic reference libraries (Collins et al. 2021) and the development of further taxon-specific assays, which may in the future accurately distinguish between the most closely related species.

Another advantage of bulk metabarcoding of processing by-products includes the ability to detect trace DNA in situations where the original tissue source is no longer available, either due to the complexity of trading operations or as a result of deliberate concealment (Challender et al. 2015). This may also allow for coarse estimation of relative volumes traded, which would be impossible through the pain-staking tissue sampling from individual specimens. Finally, dust metabarcoding is also cost-effective: the collection of dry processing residues is easier than collecting and preserving tissue samples, with a much-reduced sample size being sufficient to garner estimates of full species richness (**Figure 4c**). Technically, the collection of dust residues, compared to tissue sampling, is open to environmental contamination, whereby DNA traces can be detected from species that had passed through the sampled establishment days, weeks, and potentially months earlier. Still, this “contamination” is an inherent feature of the approach, which purposely seeks to investigate the biodiversity extracted, processed, and traded through a given hub. Certainly, a formal framework will be required and agreed by key stakeholders (traders, exporters and inspectors) on how to operationally implement shark-dust while avoiding reluctance. Possible steps include asking exporters to use brand-new containers for each batch of exports and using appropriate threshold parameters in the bioinformatic workflow.

Recent developments in fast and portable technologies open up new opportunities to run metabarcoding in the field. Our existing approach relies on laboratory equipment, which may be prohibitive in some contexts, especially in developing countries. Optimisation of third-generation sequencing technologies (Johri et al. 2019) will most likely advance *in situ* bulk metabarcoding techniques, enabling a wide range of applications in wildlife forensics and fisheries management and benefiting the global conservation community.

The CITES Secretariat promotes capacity development and the transmission of information and skills between countries in order to “efficiently, reliably, and cost-effectively identify shark items in commerce” (CoP18 Doc. 21.2), including genetic procedures. With a current list of 151 species (CITES 2022a), which now include over 50 species of requiem sharks (*Carcharhinus* spp.), over 50 species between wedgefishes and guitarfishes, as well as thresher sharks, hammerheads, mantas/devil rays and freshwater stingrays, the difficulties that countries face in complying with CITES regulations have never been greater. Decades of overexploitation have devastated elasmobranch populations; but the use of trade bans will only be successful in tandem with the implementation of reliable and cost-effective monitoring tools. Our study proposes a new method in commerce traceability from the residues of shark and ray processing where original tissue material is often unavailable. Such an approach should prove momentous for shark and ray conservation, by strengthening legality and traceability to ensure sustainability of elasmobranch populations across the world, and could inspire the design of similar methods to combat a wealth of other illegal wildlife trading activities.

## Acknowledgements

We thank all collaborators of the project Building Capacity to reduce illegal trading of shark products in Indonesia, funded under the Illegal Wildlife Trade (IWT) Challenge Fund number IWT057 and the University of Salford R&E strategy funding, and based at the Ministry for Marine Affairs and Fisheries (MMAF) – Republic of Indonesia, the University of Salford (UoS), the Centre for Environment, Fisheries and Aquaculture Science (Cefas) and the Rekam Nusantara Foundation – Indonesian (REKAM). We also thank to officers of B/LPSPLs (‘Balai/Loka Pengelolaan Sumber Daya Pesisir dan Laut’; Institute for Coastal and Marine Resource Management) and Fish Quarantine for helping during field work. We thank Holly Broadhurst and Peter Shum for training and advice in different aspects of lab techniques, Owen Wangensteen for advice on bioinformatics and Samuel Browett for advice on analyses. We also thank Efin Muttaqin and staff of the Wildlife Conservation Society – Indonesian Program (WCS-IP) for their support during project initiation. In addition, APP would personally like to thank Mr. Dharmadi and Dr. Hawis Maddupa for their legacy and supporting the younger generation of scientists seeking to make an impact in conserving elasmobranchs in Indonesia and Diego Cardeñosa for inspiring discussions.

## Permit

Research permit no. 251/BRSDM/II/2020 issued by the Agency for Marine and Fisheries Research and Human Resources AMFRAD, the Ministry of Marine Affairs and Fisheries (MMAF), Republic of Indonesia.

Research ethics no. STR1819-45 issued by Science and Technology Research Ethics Panel, the University of Salford, United Kingdom.

Export permits no. 00135/SAJI/LN/PRL/IX/2021 (CITES-listed specimens) and 127/LPSPL.2/PRL.430/X/2021 (non-CITES-listed specimens) were granted under the authority of the Ministry of Marine Affairs and Fisheries (MMAF), Republic of Indonesia.

Import permit no. 609191/01-42 from the Animal and Plant Health Agency (APHA), United Kingdom.

## Author contributions

Conceptualization: APP, ADM, SM

Funding acquisition: SM, JMM

Methodology: APP, NGS, ADM, SM

Resources: ADM, SM

Investigation: APP, NGS, ADM

Formal Analysis: APP

Visualization: APP

Project Administration: APP, JMM, FA, ADM, SM

Supervision: ADM, SM

Writing—original draft: APP, ADM, SM

Writing—review & editing: JMM, FA, NGS

## Data and materials availability

Indonesia shark and ray DNA barcodes (Elas02 fragment) have been uploaded to the NCBI Short Read Archive (SRA) under BioProject accession number PRJNA850687; and are provided in table S1. Raw sequence data OTU (presence/absence), taxa, sample metadata, bioinformatics pipeline and R scripts are available at https://github.com/andhikaprima/sharkdust and archived on Dryad: https://datadryad.org/stash/share/KKqbVy1Rf9grLEpnx_3KmW3ZnZI5ZXsm-oB24BRt_z8.

## Supplementary information

Additional Supporting Information may be found in the online version of this article:

Figure S1. Sampling locations across Java Island, Indonesia. Locations are labelled with long and short codes to facilitate identification in subsequent figures.

Figure S2. General description of sequencing results; read depth (a) and taxonomy diversity (b).

Figure S3. Workflow schematic from wet laboratory activities to bioinformatics pipeline.

Figure S4. Correlation between relative reads abundance (RRA) of species from dust samples and number of individual species from tissue samples for all sampled locations.

Figure S5. Number of raw reads per sampling site used to normalize species composition and to rank the top five species.

Table S1. Filtering steps removing all MOTUs/reads originating from sequencing errors or contamination, and the respective number of reads retrieved at each stage

Table S2. List of shark species sequenced from dust samples

Table S3. The result of PERMANOVA analysis to test for compositional differences between the two types of samples, shark-dust and individual specimen tissues

Table S4. Ambiguity in species identification

Table S5. List of analysed dust samples, including sample code, date of collection, location and notes

Table S6. List of analysed tissue samples, including sample code, date of collection, location, type of product and species identification

Table S7. List of species present in the curated reference database and the respective number of sequences included per species

