## Supplement materials for "Shark-dust: High-throughput DNA sequencing of processing residues unveils widespread trade in threatened sharks and rays"

**This PDF file includes:**

Extended materials and methods

Figure S1. Sampling locations across Java Island, Indonesia. Locations are labelled with long and short codes to facilitate identification in subsequent figures.

Figure S2. General description of sequencing results; read depth (a) and taxonomy diversity (b).

Figure S3. Workflow schematic from wet laboratory activities to bioinformatics pipeline.

Figure S4. Correlation between relative reads abundance (RRA) of species from dust samples and number of individual species from tissue samples for all sampled locations.

Figure S5. Number of raw reads per sampling site used to normalize species composition and to rank the top five species.

Table S1. Filtering steps removing all MOTUs/reads originating from sequencing errors or contamination, and the respective number of reads retrieved at each stage

Table S2. List of shark species sequenced from dust samples

Table S3. The result of PERMANOVA analysis to test for compositional differences between the two types of samples, shark-dust and individual specimen tissues

Table S4. Ambiguity in species identification

Table S5. List of analysed dust samples, including sample code, date of collection, location and notes

Table S6. List of analysed tissue samples, including sample code, date of collection, location, type of product and species identification

Table S7. List of species present in the curated reference database and the respective number of sequences included per species

### Extended materials and methods

#### *Laboratory procedures*

All laboratory works conducted with a general good laboratory practice, including all working surfaces were sterilised with 50% bleach and then washed with 70% ethanol, in-between and after extracting each sample, to reduce the risk of cross-contamination. Further measures to avoid contamination included: the use of two separate clean rooms for extraction of dust and tissue, and all the dust laboratory work (from extraction to sequencing) was conducted prior to handling the tissue samples.

The Elas02 primer sets used in this experiment were arranged into 96 different combinations of forward and reverse MID tags. These PCR plates constitutes a library of 28 samples, two PCR blanks and positive control (North Atlantic beaked redfish; *Sebastes mentella*). The PCR mix recipe was as follows: A total volume of 24 µl included 12.5 µl Qiagen™ Multiplex PCR kit, 1 µl of the 5 µM pre-mixed forward and reverse primers (Macrogen™), 3 µl of a standardised amount (10-15 ng/µl) of DNA, and 7.5 µl sterile water. The PCR profile included a 15-minute initial denaturing step at 95 °C, 40 cycles at 94 °C for 1 minute, 59 °C for 30 seconds, 72 °C for 1 minute and a 5-minute final extension step at 72 °C. After PCR, each replicate was visually examined on a 1.2% agarose gel, stained with GelRed® Nucleic Acid Gel Stain. Each well received 2 µl of sample and a 100 bp ladder Invitrogen™ was included in the gel for reference. Then, the triplicates were pooled for quantifying and bead cleaning.

In this study, complementary tissue samples were collected in a variety of ways, from being fresh to being very processed. About 180 of the 579 tissue samples that were collected for this project were selected based on where the dust samples were taken. During DNA amplification with Leray-XT primer sets, samples were distributed amongst 9 PCR plates which used 288 different combinations of forward and reverse MID tags. These 9 PCR plates were divided into three (3) libraries. The PCR mix recipe was as follows: a total volume of 15 µl included 7.5 µl Qiagen™ Multiplex PCR kit, 2 µl of the 5 µM pre-mixed forward and reverse primers (Macrogen™), 2 µl of a standardised amount (15 ng/µl) of DNA, and 3.5 µl sterile water. 5-minute final extension step at 72 °C. Each library consists of 193 samples, 5 blanks and two positive controls. The library was amplified in duplicate, but these PCR replicates were not individually barcoded. The PCR results were examined visually by gel electrophoresis prior pooled into three different libraries for proceeding to the next stage.

Before library preparation (i.e. the ligation of sequencing adapters onto PCR products), a bead clean was performed to purify the pooled PCR products from dust and tissue samples separately. A left-side bead clean was performed using MAGBio HighPrep™ PCR Clean-up System beads at a 1.1 beads:pool ratio. The purified library subset was then quantified using Qubit™ broad range (BR) kit (Thermo Fisher Scientific). The success of each cleaning step was verified on an Agilent Tapestation using High Sensitivity screen tapes. The NEXTFLEX™ single index sequencing adapters for Illumina platform were ligated onto each library. These adapters have a single 6 bp index. While libraries of tissue samples were used, three (3) unique adapter indices were associated with each library, allowing the 579 samples to be multiplexed into a sequencing run. To verify if adapters have been successfully ligated and no un-ligated adapters remain, each library was examined on the Agilent™ TapeStation using the High Sensitivity screen tapes.

#### *Enriching the reference database*

Preliminary bioinformatics analyses of the dust samples found the existing 12S marker sequence database had significant gaps and limited resolution to identify several species such as hammerhead sharks (*Sphyrna* spp.) and wedgefishes (*Rhynchobatus* spp.). To overcome this hindrance, 94 samples representing 45 species were chosen (using prior information from 650 bp of COI data; Prasetyo *et al.*, *unpublished data*) and successfully amplified using the Elas02 primer set (see protocol above). The process of PCR, bead cleaning, quantifying, adapter ligation and sequencing of reference samples followed a similar protocol for sequencing the dust samples. This library was sequenced using a MiSeq 2×150 bp nano v2 kit and was loaded at a concentration of 9 pM with a 1% PhiX spike-in 700 µl total volume. For the purpose of this study, these new sequences were added to the 12S elasmobranch database, which was last updated in July 2020 (**Table S7)**.

#### *Bioinformatics and statistical analysis*

FastQC was used to quality check reads and determine suitable length trimming. Reads were then trimmed, merged, and individual samples demultiplexed based on their unique MID tags (8 bp). Identical sequences were then collapsed before de novo detection and removal of chimaeras using VSEARCH (Rognes et al. 2016) with a minimum threshold (minh) by 0.90. We performed clustering with the default parameters of Swarm v3 (Mahé et al. 2021) with a local clustering threshold (d) at 1 and assigned the resultant sequences to taxa with ecotag and a manually curated 12S modified database. Following the pipeline, we applied strict filtering steps, that included retention of sequences within the expected size range (140 bp to 190 bp); removal of non-elasmobranch MOTUs (molecular operational taxonomic units); removal of MOTUs with a taxonomic identity of less than 97%. A minimum of two reads was required for the presence of a MOTU in a sample. Any remaining taxa that could not be assigned to phylum level in our, mostly, elasmobranch databases were manually searched in the NCBI nucleotide database using blastn and were retained if identity was greater than 97%. The read abundance of 28 samples was pooled into 7 locations where they were taken to be compared with the identification using individual tissue samples. Sequences from tissue samples using the Leray-XT primer (COI region) retained the fragment sizes between 299 bp and 320 bp and followed similar parameters to the rest. Sample identification was assigned based on the highest number of reads in an individual sample.

### Supplementary figures

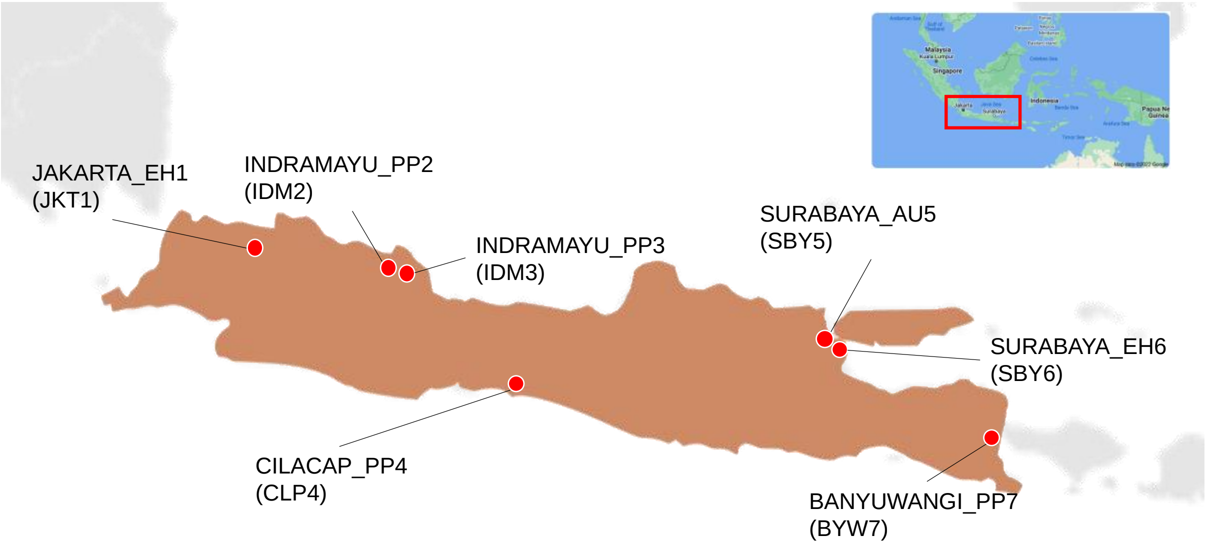


**Figure S1.** Sampling locations across Java Island, Indonesia. Locations are labelled with long and short codes to facilitate identification in subsequent figures.

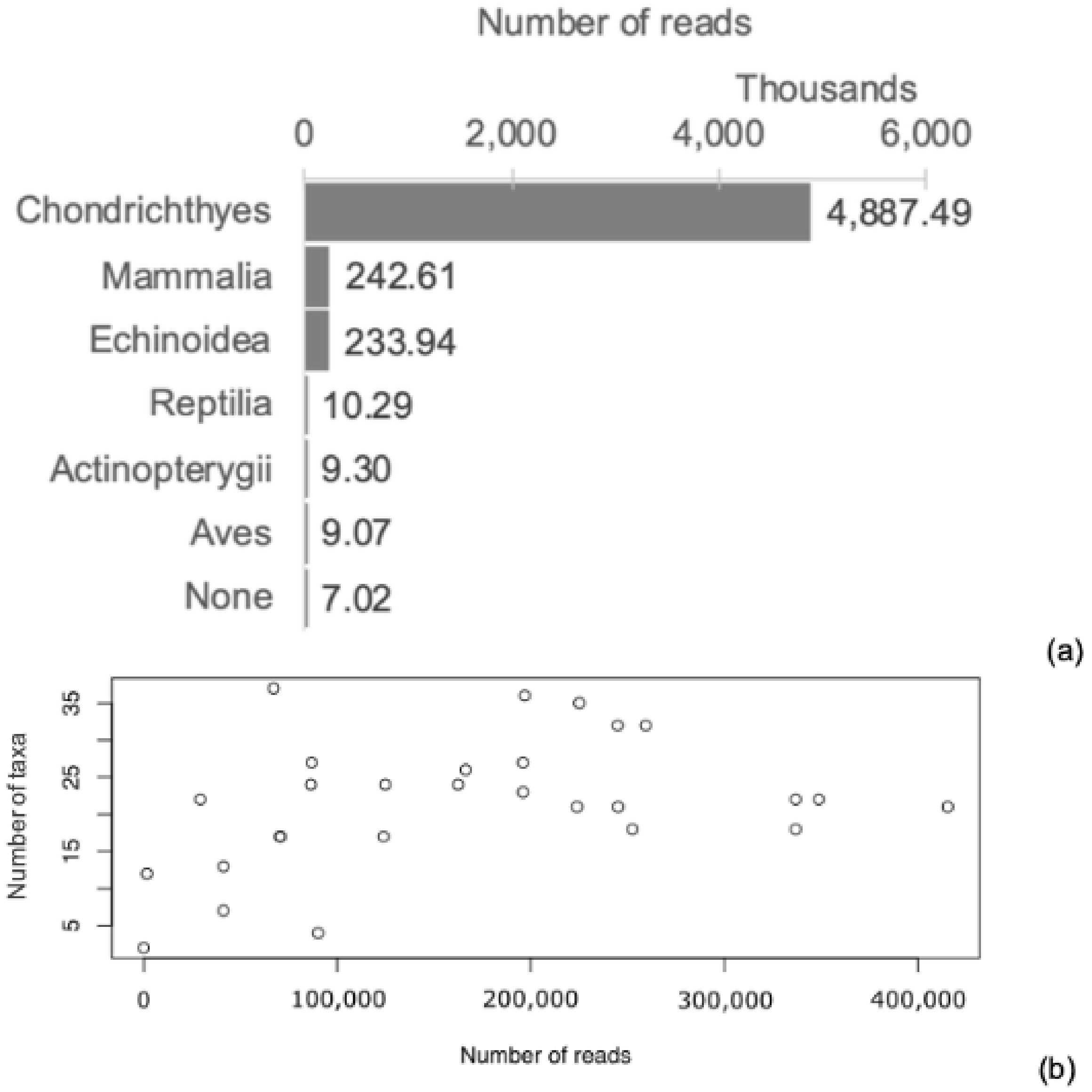


**Figure S2.** General description of sequencing results; read proportions (a) and taxonomy diversity against read numbers (b).

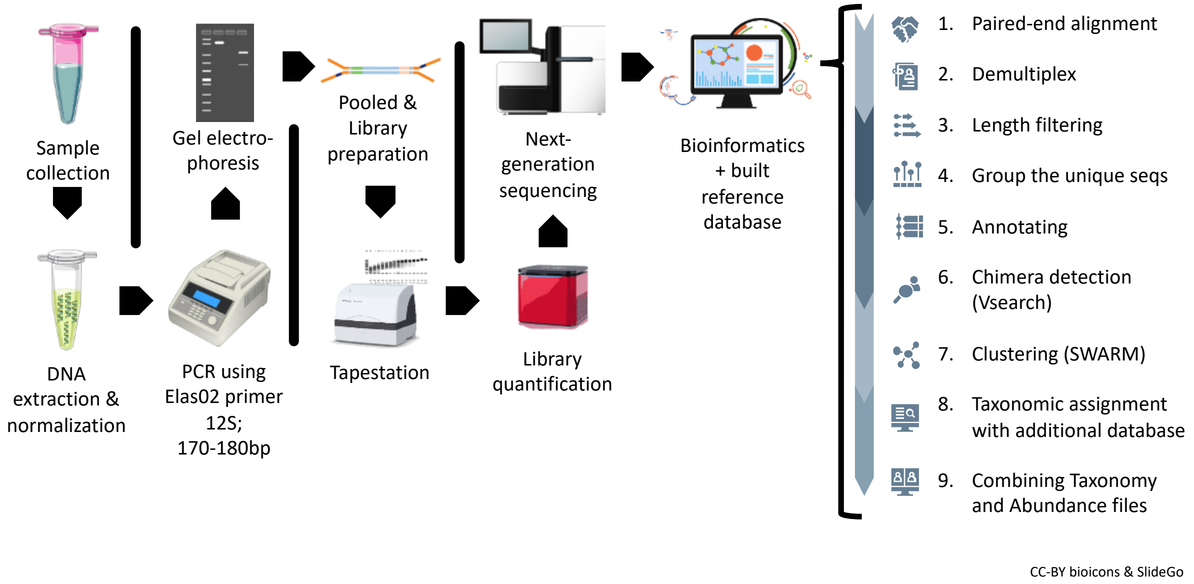


**Figure S3.** Workflow schematic from wet laboratory activities to bioinformatics pipeline of dust metabarcoding.

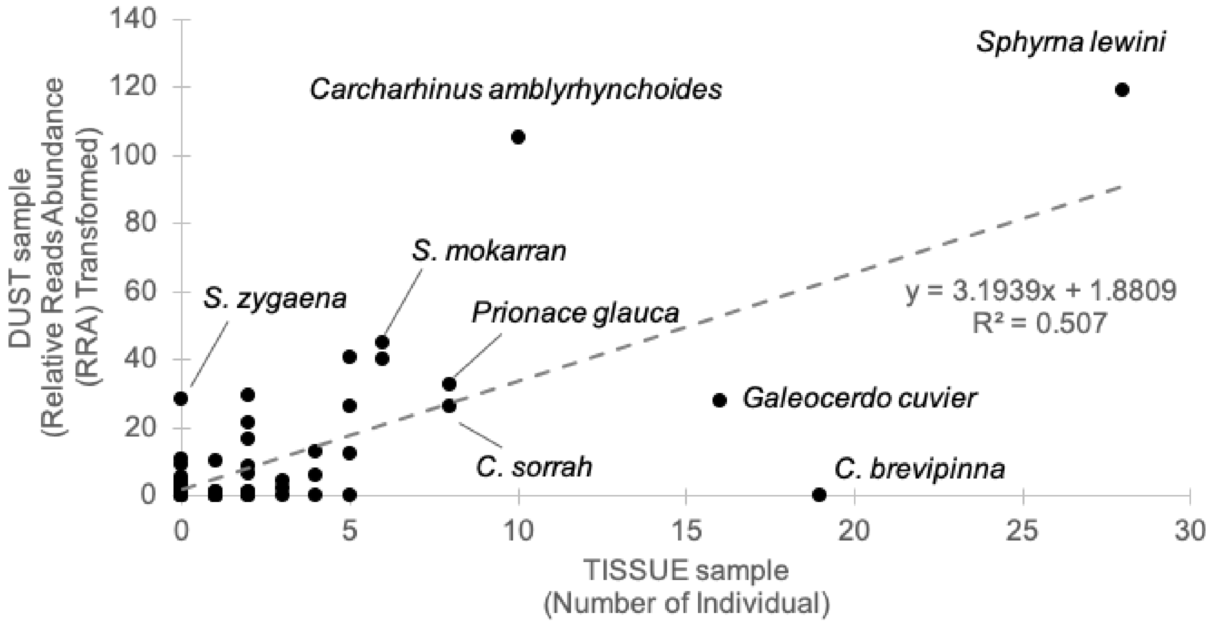


**Figure S4.** Correlation between relative reads abundance (RRA) of species from dust samples and number of individual species from tissue samples for all sampled locations.

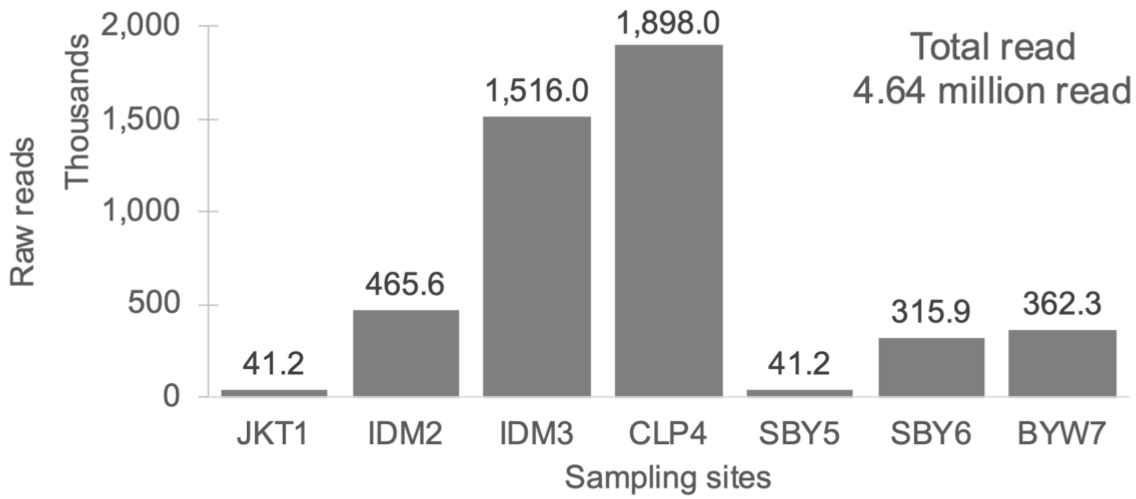


**Figure S5.** Number of raw reads per sampling site used to normalize species composition and to rank the top five species.

### Supplementary tables

**Table S1.** Filtering steps removing all MOTUs/reads originating from sequencing errors or contamination and the respective number of reads retrieved at each stage

| Filtering Steps | Total |
|---|---|
| **Total Reads** | **5,580,616** |
| After removing reads from the blanks and control | 5,098,807 |
| After removing all non-elasmobranch reads | 4,640,239 |

**Table S2.** List of shark species sequenced from dust sample and tissue sample

| Family Name | Scientific Name | English Name | Indonesian Name | CITES Status | Dust detection | Tissue detection | NCBI Accession Code |
|---|---|---|---|---|---|---|---|
| Carcharhinidae | *Prionace glauca* | Blue shark | Hiu selendang | CITES | X | X | XXX |
| Carcharhinidae | *Carcharhinus falciformis* | Silky shark | Hiu sutra | CITES | X | X | |
| Carcharhinidae | *Carcharhinus albimarginatus* | Silvertip shark | Hiu silvertip | CITES | X | X | |
| Carcharhinidae | *Carcharhinus brachyurus* | Copper shark | Hiu lanjaman | CITES | | X | |
| Carcharhinidae | *Carcharhinus brevipinna* | Spinner shark | Hiu plen | CITES | | X | |
| Carcharhinidae | *Carcharhinus longimanus* | Oceanic whitetip shark | Hiu koboi | CITES | X | X | |
| Carcharhinidae | *Carcharhinus obscurus* | Dusky shark | Hiu lanjaman | CITES | | X | |
| Carcharhinidae | *Carcharhinus plumbeus* | Sandbar shark | Hiu teteri | CITES | X | X | |
| Carcharhinidae | *Carcharhinus amblyrhynchoides* | Graceful shark | Hiu lanjaman | CITES | X | X | |
| Carcharhinidae | *Carcharhinus melanopterus* | Blacktip reef shark | Hiu mada | CITES | X | X | |
| Carcharhinidae | *Carcharhinus sorrah* | Spot-tail shark | Hiu lanjaman | CITES | X | X | |
| Carcharhinidae | *Carcharhinus leucas* | Bull shark | Hiu buas | CITES | X | X | |
| Carcharhinidae | *Carcharhinus amboinensis* | Java shark | Hiu lanjaman | CITES | X | | |
| Carcharhinidae | *Carcharhinus macloti* | Hardnose shark | Hiu aron | CITES | X | X | |
| Carcharhinidae | *Triaenodon obesus* | Whitetip reef shark | Hiu bokem | CITES | | X | |
| Carcharhinidae | *Carcharhinus dussumieri* | Whitecheek shark | Hiu lanjaman | CITES | | X | |
| Carcharhinidae | *Carcharhinus tjutjot* | Indonesian whaler shark | Hiu lanjaman | CITES | X | | |

| Family Name | Scientific Name | English Name | Indonesian Name | CITES Status | Dust detection | Tissue detection | NCBI Accession Code |
|---|---|---|---|---|---|---|---|
| Carcharhinidae | *Glyphis glyphis* | Speartooth shark | | CITES | X | | |
| Carcharhinidae | *Lamiopsis tephrodes* | Borneo broadfin shark | Hiu bujit | CITES | X | | |
| Carcharhinidae | *Scoliodon macrorhynchos* | Pacific spadenose shark | Hiu kejen | CITES | X | | |
| Carcharhinidae | *Loxodon macrorhinus* | Sliteye shark | Hiu kejen | CITES | X | | |
| Carcharhinidae | *Rhizoprionodon oligolinx* | Grey sharpnose shark | Hiu plen | CITES | X | | |
| Carcharhinidae | *Rhizoprionodon taylori* | Australian sharpnose shark | Hiu plen | CITES | | X | |
| Carcharhinidae | *Galeocerdo cuvier* | Tiger shark | Hiu macan | Non-CITES | X | X | |
| Sphyrnidae | *Eusphyra blochii* | Winghead shark | Hiu caping | CITES | X | | |
| Sphyrnidae | *Sphyrna mokarran* | Great hammerhead | Hiu caping | CITES | X | X | |
| Sphyrnidae | *Sphyrna lewini* | Scalloped hammerhead | Hiu caping | CITES | X | X | |
| Sphyrnidae | *Sphyrna zygaena* | Smooth hammerhead | Hiu caping | CITES | X | | |
| Alopiidae | *Isurus oxyrinchus* | Shortfin mako shark | Hiu tenggiri | CITES | X | X | |
| Alopiidae | *Isurus paucus* | Longfin mako shark | Hiu tenggiri | CITES | X | X | |
| Alopiidae | *Lamna nasus* | Porbeagle shark | | CITES | | X | |
| Alopiidae | *Alopias pelagicus* | Pelagic thresher | Hiu monyet | CITES | X | X | |
| Alopiidae | *Alopias superciliosus* | Bigeye thresher | Hiu monyet | CITES | X | X | |
| Hemigaleidae | *Hemigaleus australiensis* | Australian weasel shark | Hiu kacang | Non-CITES | X | X | |
| Hemigaleidae | *Hemigaleus microstoma* | Sicklefin weasel shark | Hiu kacang | Non-CITES | X | | |
| Hemigaleidae | *Hemipristis elongata* | Snaggletooth shark | Hiu monas | Non-CITES | X | | |
| Hemiscylliidae | *Chiloscyllium plagiosum* | Whitespotted bamboo | Hiu bongo | Non-CITES | X | | |

| Family Name | Scientific Name | English Name | Indonesian Name | CITES Status | Dust detection | Tissue detection | NCBI Accession Code |
|---|---|---|---|---|---|---|---|
| Hemiscylliidae | *Chiloscyllium punctatum* | Brownbanded bamboo | Hiu bongo | Non-CITES | X | X | |
| Triakidae | *Mustelus griseus* | Spotless smooth-hound | Hiu kacang | Non-CITES | X | | |
| Triakidae | *Mustelus manazo* | Starspotted smooth-hound | Hiu kacang | Non-CITES | X | X | |
| Odontaspididae | *Carcharias taurus* | Sand tiger shark | Hiu anjing | Non-CITES | X | | |
| Hexanchidae | *Hexanchus griseus* | Bluntnose sixgill shark | Hiu areuy | Non-CITES | X | | |
| Squalidae | *Squalus hemipinnis* | Indonesian shortsnout spurdog | Hiu botol | Non-CITES | X | | |
| Stegostomatidae | *Stegostoma fasciatum* | Zebra shark | Hiu belimbing | Non-CITES | X | X | |
| Dasyatidae | *Himantura gerrardi* | Whitespotted whipray | Pari bintang | Non-CITES | X | | |
| Dasyatidae | *Himantura leoparda* | Leopard whipray | Pari macan | Non-CITES | X | | |
| Dasyatidae | *Himantura uarnak* | Reticulate whipray | Pari macan | Non-CITES | | X | |
| Dasyatidae | *Pateobatis fai* | Pink whipray | Pari minyak | Non-CITES | | X | |
| Dasyatidae | *Himantura jenkinsii* | Jenkins whipray | Pari duri | Non-CITES | X | | |
| Dasyatidae | *Himantura hortlei* | Hortle's whipray | | Non-CITES | X | | |
| Dasyatidae | *Himantura granulata* | Mangrove whipray | Pari sapi | Non-CITES | X | | |
| Dasyatidae | *Urogymnus granulatus* | Mangrove whipray | | Non-CITES | | X | |
| Dasyatidae | *Neotrygon kuhlii* | Bluespotted stingray | Pari blentik | Non-CITES | X | | |
| Dasyatidae | *Dasyatis thetidis* | Thorntail stingray | | Non-CITES | X | | |
| Dasyatidae | *Dasyatis zugei* | Pale-edged stingray | Pari biasa | Non-CITES | X | | |
| Dasyatidae | *Pastinachus atrus* | Cowtail stingray | | Non-CITES | X | | |
| Myliobatidae | *Mobula birostris* | Giant oceanic manta ray | Pari kerbau | CITES | X | X | |
| Myliobatidae | *Mobula tarapacana* | Sicklefin devil ray | Pari lampingan | CITES | X | X | |
| Myliobatidae | *Mobula thurstoni* | Bentfin devil ray | Pari lampingan | CITES | X | | |

| Family Name | Scientific Name | English Name | Indonesian Name | CITES Status | Dust detection | Tissue detection | NCBI Accession Code |
|---|---|---|---|---|---|---|---|
| Myliobatidae | *Mobula mobular* | Giant devil ray | Pari lampingan | CITES | X | X | |
| Rhynchobatidae | *Rhynchobatus australiae* | Whitespotted guitarfish | Liongbun | CITES | X | X | |
| Rhynchobatidae | *Rhynchobatus laevis* | Smoothnose wedgefish | Liongbun | CITES | X | X | |
| Rhynchobatidae | *Rhynchobatus springeri* | Broadnose wedgefish | Liongbun | CITES | X | X | |
| Rhinidae | *Rhina ancylostoma* | Bowmouth guitarfish | Hiu barong | CITES | X | X | |
| Rhinobatidae | *Glaucostegus typus* | Giant guitarfish | Pari kekeh | CITES | X | X | |
| Pristidae | *Anoxypristis cuspidata* | Knifetooth sawfish | Pari gergaji lancip | CITES | X | | |
| Rhinobatidae | *Rhinobatos penggali* | Indonesian shovelnose ray | Pari kekeh | CITES | | X | |
| Gymnuridae | *Gymnura poecilura* | Longtail butterfly ray | Pari kalelawar | Non-CITES | X | | |
| Carcharhinidae | *Carcharhinus* sp. | Requiem sharks | | CITES | | | |
| Dasyatidae | *Himantura* sp. | Whiprays | | | | | |
| Myliobatidae | *Mobula* sp. | Manta/Devil rays | | CITES | | | |
| Rhinopteridae | *Rhinoptera* sp. | Cownose rays | | | | | |
| Rhynchobatidae | *Rhynchobatus* sp. | Guitarfishes | | CITES | | | |
| Carcharhinidae | | Requiem shark families | | | | | |
| Rhinobatinae | | Guitarfish families | | CITES | | | |

**Table S3.** The result of PERMANOVA analysis to test for compositional differences between the two types of samples, shark-dust and individual specimen tissues.

Permutation: free

Number of permutations: 999

| | df | Sum | MS | F.Model | $R^2$ | Pr(>F) |
|---|---|---|---|---|---|---|
| **Type** | 1 | 0.7860 | 0.78600 | 3.4976 | 0.22569 | 0.001 |
| **Residuals** | 12 | 2.6967 | 0.22472 | | 0.77431 | |
| **Total** | 13 | 3.4827 | 1.00000 | | | |

**Table S4.** Ambiguity in species identification

| Genus | Species list |
|---|---|
| 11 Carcharhinus haplotypes that could not be unambiguously assigned to one species. | *Carcharhinus amboinensis* and *Carcharhinus obscurus* |
| | *Carcharhinus plumbeus and Carcharhinus albimarginatus* |
| | *Carcharhinus amblyrhynchoides* and *Carcharhinus sorrah* |
| | *Carcharhinus falciformis, Carcharhinus amblyrhynchoides* and *Carcharhinus sorrah* |
| | *Carcharhinus acronotus, Carcharhinus porosus, Carcharhinus amboinensis* and *Carcharhinus obscurus* |
| | *Carcharhinus acronotus, Carcharhinus porosus, Carcharhinus obscurus, Carcharhinus amboinensis* and *Carcharhinus macloti* |
| | *Carcharhinus plumbeus, Carcharhinus acronotus, Carcharhinus porosus, Carcharhinus amboinensis* and *Carcharhinus obscurus* |
| | *Carcharhinus longimanus, Carcharhinus porosus, Carcharhinus obscurus, Carcharhinus amboinensis* and *Carcharhinus acronotus* |
| | *Carcharhinus acronotus, Carcharhinus porosus, Carcharhinus obscurus, Carcharhinus amboinensis* and *Carcharhinus amblyrhynchoides* |
| | *Carcharhinus plumbeus, Carcharhinus albimarginatus, Carcharhinus porosus, Carcharhinus acronotus* and *Carcharhinus amblyrhynchoides* |
| | *Carcharhinus porosus, Carcharhinus amblyrhynchoides, Carcharhinus tjutjot, Carcharhinus amboinensis, Carcharhinus acronotus* and *Carcharhinus obscurus* |
| Some genus *Himantura* | *Himantura leoparda* and *H. uarnak* |
| Some genus *Mobula* | *Mobula formosana, Mobula japanica* and *Mobula mobular* |
| | *Mobula eregoodootenkee, Mobula kuhlii* and *Mobula thurstoni* |
| Some genus *Rhinoptera* | *Rhinoptera javanica* and *R. steindachneri* |
| Some genus *Rhynchobatus* | *Rhynchobatus laevis* and *Rhynchobatus australiae* |
| | *Rhynchobatus springeri* and *Rhynchobatus djiddensis* |
| | *Rhynchobatus laevis, Rhynchobatus australiae* and *Rhynchobatus djiddensis* |

| Genus | Species list |
| --- | --- |
| Some family Carcharhinidae | *Prionace glauca, Carcharhinus acronotus* and *Carcharhinus obscurus* |
| | *Carcharhinus plumbeus, Carcharhinus porosus, Carcharhinus amblyrhynchoides, Glyphis siamensis, Glyphis fowlerae, Glyphis gangeticus, Carcharhinus leucas, Glyphis sp. Pakistan, Carcharhinus albimarginatus, Carcharhinus acronotus* and *Carcharhinus obscurus* |
| | *Carcharhinus porosus, Carcharhinus acronotus, Carcharhinus amblyrhynchoides, Carcharhinus tjutjot, Carcharhinus amboinensis, Lamiopsis temminckii* and *Carcharhinus obscurus* |
| | *Carcharhinus acronotus* and *Prionace glauca* |
| | *Carcharhinus porosus, Carcharhinus acronotus, Carcharhinus amblyrhynchoides, Carcharhinus amboinensis, Lamiopsis temminckii* and *Carcharhinus obscurus* |
| Some subfamily Rhinobatinae | *Glaucostegus formosensis, Rhinobatos schlegelii* and *Rhinobatos hynnicephalus* |

**Table S5.** List of analysed dust samples, including sample code, date of collection, location and notes

| No. | Ind ID | Pooled ID | Date | Location | Trader | Association | Notes |
|---|---|---|---|---|---|---|---|
| 1 | MB-01 | JKT1 | 9/1/20 | Muara Baru | Export hub warehouse | Fin sack | |
| 2 | IM-02 | IDM2 | 12/1/20 | Indramayu | Processing plant/collector | Fin sack | |
| 3 | IM-03 | IDM2 | 12/1/20 | Indramayu | Processing plant/collector | Fin sack | |
| 4 | IM-04 | IDM2 | 12/1/20 | Indramayu | Processing plant/collector | Fin sack | |
| 5 | IM-05 | IDM2 | 12/1/20 | Indramayu | Processing plant/collector | Fin sack | |
| 6 | IM-06 | IDM2 | 12/1/20 | Indramayu | Processing plant/collector | Fin sack | |
| 7 | IM-07 | IDM3 | 13/1/20 | Indramayu | Processing plant/collector | Fin sack | |
| 8 | IM-08 | IDM3 | 13/1/20 | Indramayu | Processing plant/collector | Fin sack | |
| 9 | IM-09 | IDM3 | 13/1/20 | Indramayu | Processing plant/collector | Fin sack | |
| 10 | IM-10 | IDM3 | 13/1/20 | Indramayu | Processing plant/collector | Fin sack | |
| 11 | IM-11 | IDM3 | 13/1/20 | Indramayu | Processing plant/collector | Cartilage sack | |
| 12 | IM-12 | IDM3 | 13/1/20 | Indramayu | Processing plant/collector | Cartilage sack | |
| 13 | IM-13 | IDM3 | 13/1/20 | Indramayu | Processing plant/collector | Cartilage sack | |
| 14 | IM-14 | IDM3 | 13/1/20 | Indramayu | Processing plant/collector | Cartilage sack | |
| 15 | IM-15 | IDM3 | 13/1/20 | Indramayu | Processing plant/collector | Skin pile | |
| 16 | IM-16 | IDM3 | 13/1/20 | Indramayu | Processing plant/collector | Skin pile | |
| 17 | IM-17 | IDM3 | 13/1/20 | Indramayu | Processing plant/collector | Skin pile | Not enough sample quantity |
| 18 | IM-18 | IDM3 | 13/1/20 | Indramayu | Processing plant/collector | Skin pile | Not enough sample quantity |
| 19 | IM-19 | IDM3 | 13/1/20 | Indramayu | Processing plant/collector | Meat boxes | Not enough sample quantity |
| 20 | CL-20 | CLP4 | 25/1/20 | Cilacap | Processing plant/collector | Fin sack | |
| 21 | CL-21 | CLP4 | 25/1/20 | Cilacap | Processing plant/collector | Fin dust from saw machine | |
| 22 | CL-22 | CLP4 | 25/1/20 | Cilacap | Processing plant/collector | Fin dust from saw machine | |
| 23 | CL-23 | CLP4 | 25/1/20 | Cilacap | Processing plant/collector | Fin dust from saw machine | |
| 24 | CL-24 | CLP4 | 25/1/20 | Cilacap | Processing plant/collector | Fin dust from saw machine | |

| No. | Ind ID | Pooled ID | Date | Location | Trader | Association | Notes |
|---|---|---|---|---|---|---|---|
| 25 | CL-25 | CLP4 | 26/1/20 | Cilacap | Processing plant/collector | Drying places for meat, skin, cartilage and other fishes | |
| 26 | CL-26 | CLP4 | 26/1/20 | Cilacap | Processing plant/collector | Drying places for meat, skin, cartilage and other fishes | |
| 27 | SB-27 | SBY5 | 28/1/20 | Surabaya | Authority | Products collection | |
| 28 | SB-28 | SBY6 | 29/1/20 | Surabaya | Export hub warehouse | Fin sack | |
| 29 | SB-29 | SBY6 | 29/1/20 | Surabaya | Export hub warehouse | Fin sack | |
| 30 | BW-30 | BYW7 | 2/2/20 | Banyuwangi | Processing plant/collector | Drying places for skin, cartilage and lower lobe caudal fin in PPP Muncar | |
| 31 | BW-31 | BYW7 | 2/2/20 | Banyuwangi | Processing plant/collector | Drying places for skin, cartilage and lower lobe caudal fin in PPP Muncar | |

Notes: Processing plants (PP), export hubs (EH) and an inspector station (AU)

**Table S6.** List of analysed tissue samples, including sample code, date of collection, location, type of product and species identification

| No. | ID | Date | Location | Dust Pooled ID Location | Type of Location | Type of Product | Part | Species Identification | CITES Status |
|---|---|---|---|---|---|---|---|---|---|
| 1 | MB-50 | 9/1/20 | Muara Baru | JKT1 | EH | Processed | Dried fin | *Isurus oxyrinchus* | CITES |
| 2 | MB-51 | 9/1/20 | Muara Baru | JKT1 | EH | Processed | Dried fin | *Lamna nasus* | CITES |
| 3 | MB-52 | 9/1/20 | Muara Baru | JKT1 | EH | Processed | Dried fin | *Isurus paucus* | CITES |
| 4 | MB-53 | 9/1/20 | Muara Baru | JKT1 | EH | Processed | Dried fin | *Carcharhinus longimanus* | CITES |
| 5 | MB-54 | 9/1/20 | Muara Baru | JKT1 | EH | Processed | Dried fin | *Alopias superciliosus* | CITES |
| 6 | IM-111 | 12/1/20 | Indramayu | IDM2 | PP | Processed | Dried fin | Unidentified | |
| 7 | IM-112 | 12/1/20 | Indramayu | IDM2 | PP | Fresh | Finless | *Sphyrna lewini* | CITES |
| 8 | IM-113 | 12/1/20 | Indramayu | IDM2 | PP | Fresh | Finless | *Sphyrna mokarran* | CITES |
| 9 | IM-114 | 12/1/20 | Indramayu | IDM2 | PP | Fresh | Finless | *Carcharhinus brevipinna* | CITES |
| 10 | IM-115 | 12/1/20 | Indramayu | IDM2 | PP | Fresh | Whole | *Sphyrna lewini* | CITES |
| 11 | IM-116 | 12/1/20 | Indramayu | IDM2 | PP | Fresh | Whole | *Carcharhinus sorrah* | CITES |
| 12 | IM-117 | 12/1/20 | Indramayu | IDM2 | PP | Fresh | Whole | *Carcharhinus sorrah* | CITES |
| 13 | IM-118 | 12/1/20 | Indramayu | IDM2 | PP | Fresh | Whole | *Hemigaleus australiensis* | Non-CITES |
| 14 | IM-119 | 12/1/20 | Indramayu | IDM2 | PP | Fresh | Whole | *Carcharhinus macloti* | CITES |
| 15 | IM-120 | 12/1/20 | Indramayu | IDM2 | PP | Fresh | Whole | *Carcharhinus amblyrhynchoides* | CITES |
| 16 | IM-121 | 12/1/20 | Indramayu | IDM2 | PP | Fresh | Whole | *Sphyrna lewini* | CITES |
| 17 | IM-122 | 12/1/20 | Indramayu | IDM2 | PP | Fresh | Finless | *Sphyrna lewini* | CITES |
| 18 | IM-123 | 12/1/20 | Indramayu | IDM2 | PP | Fresh | Finless | *Carcharhinus brevipinna* | CITES |
| 19 | IM-124 | 12/1/20 | Indramayu | IDM2 | PP | Fresh | Finless | *Carcharhinus amblyrhynchoides* | CITES |
| 20 | IM-125 | 12/1/20 | Indramayu | IDM2 | PP | Fresh | Finless | *Sphyrna lewini* | CITES |
| 21 | IM-126 | 12/1/20 | Indramayu | IDM2 | PP | Fresh | Finless | *Sphyrna lewini* | CITES |
| 22 | IM-127 | 12/1/20 | Indramayu | IDM2 | PP | Fresh | Whole | *Carcharhinus sorrah* | CITES |
| 23 | IM-128 | 12/1/20 | Indramayu | IDM2 | PP | Fresh | Whole | *Carcharhinus sorrah* | CITES |

| No. | ID | Date | Location | Dust Pooled ID Location | Type of Location | Type of Product | Part | Species Identification | CITES Status |
|---|---|---|---|---|---|---|---|---|---|
| 24 | IM-129 | 12/1/20 | Indramayu | IDM2 | PP | Fresh | Whole | *Carcharhinus amblyrhynchoides* | CITES |
| 25 | IM-130 | 12/1/20 | Indramayu | IDM2 | PP | Fresh | Finless | *Carcharhinus amblyrhynchoides* | CITES |
| 26 | IM-131 | 12/1/20 | Indramayu | IDM2 | PP | Fresh | Finless | *Carcharhinus amblyrhynchoides* | CITES |
| 27 | IM-132 | 12/1/20 | Indramayu | IDM2 | PP | Fresh | Whole | *Hemigaleus australiensis* | Non-CITES |
| 28 | IM-177 | 13/1/20 | Indramayu | IDM3 | PP | Fresh | Finless | *Rhynchobatus laevis* | CITES |
| 29 | IM-178 | 13/1/20 | Indramayu | IDM3 | PP | Fresh | Whole | *Galeocerdo cuvier* | Non-CITES |
| 30 | IM-179 | 13/1/20 | Indramayu | IDM3 | PP | Fresh | Trunk | *Stegostoma fasciatum* | Non-CITES |
| 31 | IM-180 | 13/1/20 | Indramayu | IDM3 | PP | Fresh | Trunk | *Stegostoma fasciatum* | Non-CITES |
| 32 | IM-181 | 13/1/20 | Indramayu | IDM3 | PP | Fresh | Trunk | *Stegostoma fasciatum* | Non-CITES |
| 33 | IM-182 | 13/1/20 | Indramayu | IDM3 | PP | Fresh | Finless | *Carcharhinus longimanus* | CITES |
| 34 | IM-183 | 13/1/20 | Indramayu | IDM3 | PP | Fresh | Whole | *Carcharhinus amblyrhynchoides* | CITES |
| 35 | IM-184 | 13/1/20 | Indramayu | IDM3 | PP | Fresh | Trunk | *Sphyrna lewini* | CITES |
| 36 | IM-185 | 13/1/20 | Indramayu | IDM3 | PP | Fresh | Whole | *Carcharhinus sorrah* | CITES |
| 37 | IM-186 | 13/1/20 | Indramayu | IDM3 | PP | Fresh | Trunk | *Sphyrna lewini* | CITES |
| 38 | IM-187 | 13/1/20 | Indramayu | IDM3 | PP | Fresh | Trunk | *Sphyrna lewini* | CITES |
| 39 | IM-188 | 14/1/20 | Indramayu | IDM3 | PP | Fresh | Whole | *Carcharhinus sorrah* | CITES |
| 40 | IM-189 | 14/1/20 | Indramayu | IDM3 | PP | Fresh | Whole | *Rhynchobatus springeri* | CITES |
| 41 | IM-190 | 14/1/20 | Indramayu | IDM3 | PP | Fresh | Whole | *Hemigaleus australiensis* | Non-CITES |
| 42 | IM-191 | 13/1/20 | Indramayu | IDM3 | PP | Processed | Whole Salted | *Chiloscyllium punctatum* | Non-CITES |
| 43 | IM-192 | 13/1/20 | Indramayu | IDM3 | PP | Processed | Whole Salted | *Rhizoprionodon taylori* | CITES |
| 44 | IM-193 | 13/1/20 | Indramayu | IDM3 | PP | Processed | Cartillage | Unidentified | |
| 45 | IM-194 | 13/1/20 | Indramayu | IDM3 | PP | Processed | Cartillage | *Sphyrna lewini* | CITES |
| 46 | IM-195 | 13/1/20 | Indramayu | IDM3 | PP | Processed | Dried skin | *Stegostoma fasciatum* | Non-CITES |
| 47 | IM-196 | 13/1/20 | Indramayu | IDM3 | PP | Processed | Dried skin | *Glaucostegus typus* | CITES |

| No. | ID | Date | Location | Dust Pooled ID Location | Type of Location | Type of Product | Part | Species Identification | CITES Status |
|---|---|---|---|---|---|---|---|---|---|
| 48 | IM-197 | 13/1/20 | Indramayu | IDM3 | PP | Processed | Dried skin | *Sphyrna mokarran* | CITES |
| 49 | CL-338 | 25/1/20 | Cilacap | CPL4 | PP | Processed | Dried fin | *Isurus paucus* | CITES |
| 50 | CL-339 | 25/1/20 | Cilacap | CPL4 | PP | Processed | Dried fin | *Isurus paucus* | CITES |
| 51 | CL-340 | 25/1/20 | Cilacap | CPL4 | PP | Processed | Dried fin | *Alopias pelagicus* | CITES |
| 52 | CL-341 | 25/1/20 | Cilacap | CPL4 | PP | Processed | Dried fin | *Alopias pelagicus* | CITES |
| 53 | CL-341X | 25/1/20 | Cilacap | CPL4 | PP | Processed | Dried fin | *Carcharhinus longimanus* | CITES |
| 54 | CL-342 | 25/1/20 | Cilacap | CPL4 | PP | Processed | Dried fin | *Carcharhinus longimanus* | CITES |
| 55 | CL-343 | 25/1/20 | Cilacap | CPL4 | PP | Processed | Dried fin | *Isurus oxyrinchus* | CITES |
| 56 | CL-344 | 25/1/20 | Cilacap | CPL4 | PP | Processed | Dried fin | *Isurus oxyrinchus* | CITES |
| 57 | CL-345 | 25/1/20 | Cilacap | CPL4 | PP | Processed | Dried fin | *Alopias superciliosus* | CITES |
| 58 | CL-346 | 25/1/20 | Cilacap | CPL4 | PP | Processed | Dried fin | *Alopias superciliosus* | CITES |
| 59 | CL-347 | 25/1/20 | Cilacap | CPL4 | PP | Processed | Dried fin | *Carcharhinus brevipinna* | CITES |
| 60 | CL-348 | 25/1/20 | Cilacap | CPL4 | PP | Processed | Dried fin | *Carcharhinus brachyurus* | CITES |
| 61 | CL-349 | 25/1/20 | Cilacap | CPL4 | PP | Processed | Dried fin | *Carcharhinus brachyurus* | CITES |
| 62 | CL-350 | 25/1/20 | Cilacap | CPL4 | PP | Processed | Dried fin | *Carcharhinus leucas* | CITES |
| 63 | CL-351 | 25/1/20 | Cilacap | CPL4 | PP | Processed | Dried fin | *Carcharhinus plumbeus* | CITES |
| 64 | CL-352 | 25/1/20 | Cilacap | CPL4 | PP | Processed | Dried fin | *Carcharhinus leucas* | CITES |
| 65 | CL-353 | 25/1/20 | Cilacap | CPL4 | PP | Processed | Dried fin | *Carcharhinus leucas* | CITES |
| 66 | CL-354 | 25/1/20 | Cilacap | CPL4 | PP | Processed | Dried fin | *Galeocerdo cuvier* | Non-CITES |
| 67 | CL-355 | 25/1/20 | Cilacap | CPL4 | PP | Processed | Dried fin | *Prionace glauca* | CITES |
| 68 | CL-356 | 25/1/20 | Cilacap | CPL4 | PP | Processed | Dried fin | *Prionace glauca* | CITES |
| 69 | CL-357 | 25/1/20 | Cilacap | CPL4 | PP | Processed | Cartillage | *Alopias superciliosus* | CITES |
| 70 | CL-358 | 25/1/20 | Cilacap | CPL4 | PP | Processed | Cartillage | *Carcharhinus amblyrhynchoides* | CITES |
| 71 | CL-359 | 25/1/20 | Cilacap | CPL4 | PP | Processed | Dried fin | *Carcharhinus brevipinna* | CITES |
| 72 | CL-360 | 25/1/20 | Cilacap | CPL4 | PP | Processed | Dried fin | *Urogymnus granulatus* | Non-CITES |

| No. | ID | Date | Location | Dust Pooled ID Location | Type of Location | Type of Product | Part | Species Identification | CITES Status |
|---|---|---|---|---|---|---|---|---|---|
| 73 | CL-363 | 26/1/20 | Cilacap | CPL4 | PP | Fresh | Whole | *Galeocerdo cuvier* | Non-CITES |
| 74 | CL-364 | 26/1/20 | Cilacap | CPL4 | PP | Processed | Dried skin | *Sphyrna mokarran* | CITES |
| 75 | CL-365 | 26/1/20 | Cilacap | CPL4 | PP | Processed | Dried skin | *Carcharhinus brevipinna* | CITES |
| 76 | CL-366 | 26/1/20 | Cilacap | CPL4 | PP | Processed | Salted meat | *Alopias superciliosus* | CITES |
| 77 | CL-367 | 26/1/20 | Cilacap | CPL4 | PP | Processed | Salted meat | *Sphyrna mokarran* | CITES |
| 78 | CL-368 | 26/1/20 | Cilacap | CPL4 | PP | Processed | Salted meat | *Pateobatis fai* | Non-CITES |
| 79 | CL-369 | 26/1/20 | Cilacap | CPL4 | PP | Processed | Salted meat | *Hemigaleus australiensis* | Non-CITES |
| 80 | CL-370 | 26/1/20 | Cilacap | CPL4 | PP | Processed | Salted meat | *Mobula birostris* | CITES |
| 81 | CL-371 | 26/1/20 | Cilacap | CPL4 | PP | Processed | Salted meat | *Rhinobatos penggali* | CITES |
| 82 | CL-372 | 26/1/20 | Cilacap | CPL4 | PP | Processed | Salted meat | *Carcharhinus brevipinna* | CITES |
| 83 | CL-373 | 26/1/20 | Cilacap | CPL4 | PP | Processed | Salted meat | *Rhinobatos penggali* | CITES |
| 84 | CL-374 | 26/1/20 | Cilacap | CPL4 | PP | Processed | Salted meat | *Carcharhinus sorrah* | CITES |
| 85 | CL-375 | 26/1/20 | Cilacap | CPL4 | PP | Processed | Salted meat | *Rhinobatos penggali* | CITES |
| 86 | CL-376 | 26/1/20 | Cilacap | CPL4 | PP | Processed | Salted meat | *Carcharhinus amblyrhynchoides* | CITES |
| 87 | CL-377 | 26/1/20 | Cilacap | CPL4 | PP | Processed | Salted meat | *Carcharhinus falciformis* | CITES |
| 88 | CL-378 | 26/1/20 | Cilacap | CPL4 | PP | Processed | Salted meat | *Himantura uarnak* | Non-CITES |
| 89 | CL-380 | 26/1/20 | Cilacap | CPL4 | PP | Processed | Salted meat | *Pateobatis fai* | Non-CITES |

| No. | ID | Date | Location | Dust Pooled ID Location | Type of Location | Type of Product | Part | Species Identification | CITES Status |
|---|---|---|---|---|---|---|---|---|---|
| 90 | SB-381 | 28/1/20 | Surabaya | SBY5 | AU | Processed | Dried fin | *Carcharhinus sorrah* | CITES |
| 91 | SB-382 | 28/1/20 | Surabaya | SBY5 | AU | Processed | Dried fin | *Rhynchobatus springeri* | CITES |
| 92 | SB-383 | 28/1/20 | Surabaya | SBY5 | AU | Processed | Dried fin | *Rhina ancylostoma* | CITES |
| 93 | SB-384 | 28/1/20 | Surabaya | SBY5 | AU | Processed | Dried fin | *Isurus oxyrinchus* | CITES |
| 94 | SB-385 | 28/1/20 | Surabaya | SBY5 | AU | Processed | Dried fin | *Carcharhinus obscurus* | CITES |
| 95 | SB-386 | 28/1/20 | Surabaya | SBY5 | AU | Processed | Dried fin | *Carcharhinus amblyrhynchoides* | CITES |
| 96 | SB-387 | 28/1/20 | Surabaya | SBY5 | AU | Processed | Dried fin | *Carcharhinus leucas* | CITES |
| 97 | SB-388 | 28/1/20 | Surabaya | SBY5 | AU | Processed | Dried fin | *Triaenodon obesus* | CITES |
| 98 | SB-389 | 28/1/20 | Surabaya | SBY5 | AU | Processed | Dried fin | *Carcharhinus obscurus* | CITES |
| 99 | SB-391 | 28/1/20 | Surabaya | SBY5 | AU | Processed | Dried fin | *Carcharhinus albimarginatus* | CITES |
| 100 | SB-392 | 28/1/20 | Surabaya | SBY5 | AU | Processed | Cartillage | *Prionace glauca* | CITES |
| 101 | SB-393 | 28/1/20 | Surabaya | SBY5 | AU | Processed | Dried skin | *Carcharhinus dussumieri* | CITES |
| 102 | SB-394 | 28/1/20 | Surabaya | SBY5 | AU | Processed | Dried fin unskin | *Carcharhinus macloti* | CITES |
| 103 | SB-395 | 28/1/20 | Surabaya | SBY5 | AU | Processed | Oil | Unidentified | |
| 104 | SB-396 | 28/1/20 | Surabaya | SBY5 | AU | Processed | Oil | Unidentified | |
| 105 | SB-397 | 28/1/20 | Surabaya | SBY5 | AU | Processed | Cartillage powder | *Mobula tarapacana* | CITES |
| 106 | SB-398 | 28/1/20 | Surabaya | SBY5 | AU | Processed | Cartillage fin | *Carcharhinus brevipinna* | CITES |
| 107 | SB-399 | 28/1/20 | Surabaya | SBY5 | AU | Processed | Dried fin unskin | *Prionace glauca* | CITES |
| 108 | SB-400 | 28/1/20 | Surabaya | SBY5 | AU | Processed | Dried fin unskin | *Prionace glauca* | CITES |
| 109 | SB-401 | 28/1/20 | Surabaya | SBY5 | AU | Processed | Dried fin hissit | *Sphyrna lewini* | CITES |
| 110 | SB-402 | 28/1/20 | Surabaya | SBY5 | AU | Processed | Dried fin unskin | *Carcharhinus dussumieri* | CITES |

| No. | ID | Date | Location | Dust Pooled ID Location | Type of Location | Type of Product | Part | Species Identification | CITES Status |
|---|---|---|---|---|---|---|---|---|---|
| 111 | SB-403 | 28/1/20 | Surabaya | SBY5 | AU | Processed | Cartillage fin | *Mustelus manazo* | Non-CITES |
| 112 | SB-404 | 28/1/20 | Surabaya | SBY5 | AU | Processed | Dried skin | *Carcharhinus leucas* | CITES |
| 113 | SB-405 | 28/1/20 | Surabaya | SBY5 | AU | Processed | Dried fin unskin | *Mustelus manazo* | Non-CITES |
| 114 | SB-406 | 28/1/20 | Surabaya | SBY5 | AU | Processed | Cartillage | *Prionace glauca* | CITES |
| 115 | SB-407 | 28/1/20 | Surabaya | SBY5 | AU | Processed | Cartillage powder | Unidentified | |
| 116 | SB-408 | 28/1/20 | Surabaya | SBY5 | AU | Processed | Cartillage powder | Unidentified | |
| 117 | SB-409 | 28/1/20 | Surabaya | SBY5 | AU | Processed | Cartillage powder | Unidentified | |
| 118 | SB-410 | 28/1/20 | Surabaya | SBY5 | AU | Processed | Dried fin | *Prionace glauca* | CITES |
| 119 | SB-411 | 28/1/20 | Surabaya | SBY5 | AU | Processed | Dried fin | *Carcharhinus longimanus* | CITES |
| 120 | SB-412 | 28/1/20 | Surabaya | SBY5 | AU | Processed | Gill racker | *Mobula birostris* | CITES |
| 121 | SB-418 | 29/1/20 | Surabaya | SBY6 | EH | Processed | Dried fin | *Sphyrna mokarran* | CITES |
| 122 | SB-419 | 29/1/20 | Surabaya | SBY6 | EH | Processed | Dried fin | *Sphyrna lewini* | CITES |
| 123 | SB-420 | 29/1/20 | Surabaya | SBY6 | EH | Processed | Dried fin | *Sphyrna mokarran* | CITES |
| 124 | SB-421 | 29/1/20 | Surabaya | SBY6 | EH | Processed | Dried fin | *Isurus oxyrinchus* | CITES |
| 125 | SB-422 | 29/1/20 | Surabaya | SBY6 | EH | Processed | Dried fin | *Glaucostegus typus* | CITES |
| 126 | SB-423 | 29/1/20 | Surabaya | SBY6 | EH | Processed | Dried fin | *Rhina ancylostoma* | CITES |
| 127 | SB-424 | 29/1/20 | Surabaya | SBY6 | EH | Processed | Dried fin | *Rhynchobatus australiae* | CITES |
| 128 | SB-425 | 29/1/20 | Surabaya | SBY6 | EH | Processed | Dried fin | *Rhynchobatus springeri* | CITES |
| 129 | BW-432 | 2/2/20 | Banyuwangi | BYW7 | PP | Processed | Dried fin | *Galeocerdo cuvier* | Non-CITES |
| 130 | BW-433 | 2/2/20 | Banyuwangi | BYW7 | PP | Processed | Dried fin | *Galeocerdo cuvier* | Non-CITES |
| 131 | BW-434 | 2/2/20 | Banyuwangi | BYW7 | PP | Processed | Dried fin | *Galeocerdo cuvier* | Non-CITES |
| 132 | BW-435 | 2/2/20 | Banyuwangi | BYW7 | PP | Processed | Dried fin | *Galeocerdo cuvier* | Non-CITES |

| No. | ID | Date | Location | Dust Pooled ID Location | Type of Location | Type of Product | Part | Species Identification | CITES Status |
|---|---|---|---|---|---|---|---|---|---|
| 133 | BW-436 | 2/2/20 | Banyuwangi | BYW7 | PP | Processed | Dried fin | *Galeocerdo cuvier* | Non-CITES |
| 134 | BW-437 | 2/2/20 | Banyuwangi | BYW7 | PP | Processed | Dried fin | *Galeocerdo cuvier* | Non-CITES |
| 135 | BW-438 | 2/2/20 | Banyuwangi | BYW7 | PP | Processed | Dried fin | *Galeocerdo cuvier* | Non-CITES |
| 136 | BW-439 | 2/2/20 | Banyuwangi | BYW7 | PP | Processed | Teeth | *Galeocerdo cuvier* | Non-CITES |
| 137 | BW-440 | 2/2/20 | Banyuwangi | BYW7 | PP | Processed | Cartillage | Unidentified | |
| 138 | BW-441 | 2/2/20 | Banyuwangi | BYW7 | PP | Processed | Dried skin | *Galeocerdo cuvier* | Non-CITES |
| 139 | BW-442 | 2/2/20 | Banyuwangi | BYW7 | PP | Processed | Dried fin | *Carcharhinus amblyrhynchoides* | CITES |
| 140 | BW-443 | 2/2/20 | Banyuwangi | BYW7 | PP | Processed | Dried fin | *Sphyrna lewini* | CITES |
| 141 | BW-444 | 2/2/20 | Banyuwangi | BYW7 | PP | Processed | Dried fin | *Carcharhinus brevipinna* | CITES |
| 142 | BW-445 | 2/2/20 | Banyuwangi | BYW7 | PP | Processed | Dried fin | *Sphyrna lewini* | CITES |
| 143 | BW-446 | 2/2/20 | Banyuwangi | BYW7 | PP | Processed | Dried fin | *Sphyrna lewini* | CITES |
| 144 | BW-447 | 2/2/20 | Banyuwangi | BYW7 | PP | Processed | Dried fin | *Sphyrna lewini* | CITES |
| 145 | BW-448 | 2/2/20 | Banyuwangi | BYW7 | PP | Processed | Gill racker | *Mobula mobular* | CITES |
| 146 | BW-449 | 2/2/20 | Banyuwangi | BYW7 | PP | Processed | Gill racker | *Mobula mobular* | CITES |
| 147 | BW-450 | 2/2/20 | Banyuwangi | BYW7 | PP | Processed | Gill racker | *Mobula mobular* | CITES |
| 148 | BW-451 | 2/2/20 | Banyuwangi | BYW7 | PP | Processed | Gill racker | *Mobula mobular* | CITES |
| 149 | BW-452 | 2/2/20 | Banyuwangi | BYW7 | PP | Processed | Salted meat | *Carcharhinus melanopterus* | CITES |
| 150 | BW-452X | 2/2/20 | Banyuwangi | BYW7 | PP | Processed | Salted meat | *Carcharhinus melanopterus* | CITES |
| 151 | BW-453 | 2/2/20 | Banyuwangi | BYW7 | PP | Fresh | Fin | *Prionace glauca* | CITES |
| 152 | BW-454 | 2/2/20 | Banyuwangi | BYW7 | PP | Fresh | Fin | *Galeocerdo cuvier* | Non-CITES |
| 153 | BW-455 | 2/2/20 | Banyuwangi | BYW7 | PP | Fresh | Fin | *Galeocerdo cuvier* | Non-CITES |
| 154 | BW-456 | 2/2/20 | Banyuwangi | BYW7 | PP | Fresh | Fin | *Galeocerdo cuvier* | Non-CITES |

| No. | ID | Date | Location | Dust Pooled ID Location | Type of Location | Type of Product | Part | Species Identification | CITES Status |
|---|---|---|---|---|---|---|---|---|---|
| 155 | BW-457 | 2/2/20 | Banyuwangi | BYW7 | PP | Fresh | Fin | *Carcharhinus falciformis* | CITES |
| 156 | BW-458 | 2/2/20 | Banyuwangi | BYW7 | PP | Fresh | Fin | *Carcharhinus falciformis* | CITES |
| 157 | BW-459 | 2/2/20 | Banyuwangi | BYW7 | PP | Fresh | Fin | *Sphyrna lewini* | CITES |
| 158 | BW-460 | 2/2/20 | Banyuwangi | BYW7 | PP | Fresh | Fin | *Carcharhinus brevipinna* | CITES |
| 159 | BW-461 | 3/2/20 | Banyuwangi | BYW7 | PP | Fresh | Whole | *Sphyrna lewini* | CITES |
| 160 | BW-462 | 3/2/20 | Banyuwangi | BYW7 | PP | Fresh | Whole | *Sphyrna lewini* | CITES |
| 161 | BW-463 | 3/2/20 | Banyuwangi | BYW7 | PP | Fresh | Whole | *Carcharhinus brevipinna* | CITES |
| 162 | BW-464 | 3/2/20 | Banyuwangi | BYW7 | PP | Fresh | Whole | *Carcharhinus brevipinna* | CITES |
| 163 | BW-465 | 3/2/20 | Banyuwangi | BYW7 | PP | Fresh | Whole | *Galeocerdo cuvier* | Non-CITES |
| 164 | BW-466 | 3/2/20 | Banyuwangi | BYW7 | PP | Fresh | Finless | *Carcharhinus falciformis* | CITES |
| 165 | BW-467 | 3/2/20 | Banyuwangi | BYW7 | PP | Fresh | Whole | *Carcharhinus brevipinna* | CITES |
| 166 | BW-468 | 3/2/20 | Banyuwangi | BYW7 | PP | Fresh | Whole | *Carcharhinus brevipinna* | CITES |
| 167 | BW-469 | 3/2/20 | Banyuwangi | BYW7 | PP | Fresh | Whole | *Sphyrna lewini* | CITES |
| 168 | BW-470 | 3/2/20 | Banyuwangi | BYW7 | PP | Fresh | Whole | *Sphyrna lewini* | CITES |
| 169 | BW-471 | 3/2/20 | Banyuwangi | BYW7 | PP | Fresh | Whole | *Sphyrna lewini* | CITES |
| 170 | BW-472 | 3/2/20 | Banyuwangi | BYW7 | PP | Fresh | Whole | *Carcharhinus brevipinna* | CITES |
| 171 | BW-473 | 3/2/20 | Banyuwangi | BYW7 | PP | Fresh | Whole | *Carcharhinus brevipinna* | CITES |
| 172 | BW-474 | 3/2/20 | Banyuwangi | BYW7 | PP | Fresh | Whole | *Carcharhinus falciformis* | CITES |
| 173 | BW-475 | 3/2/20 | Banyuwangi | BYW7 | PP | Fresh | Whole | *Carcharhinus brevipinna* | CITES |
| 174 | BW-476 | 3/2/20 | Banyuwangi | BYW7 | PP | Fresh | Whole | *Sphyrna lewini* | CITES |
| 175 | BW-477 | 3/2/20 | Banyuwangi | BYW7 | PP | Fresh | Whole | *Carcharhinus brevipinna* | CITES |
| 176 | BW-478 | 3/2/20 | Banyuwangi | BYW7 | PP | Fresh | Whole | *Carcharhinus falciformis* | CITES |
| 177 | BW-479 | 3/2/20 | Banyuwangi | BYW7 | PP | Fresh | Whole | *Carcharhinus brevipinna* | CITES |
| 178 | BW-480 | 3/2/20 | Banyuwangi | BYW7 | PP | Fresh | Whole | *Carcharhinus brevipinna* | CITES |
| 179 | BW-481 | 3/2/20 | Banyuwangi | BYW7 | PP | Fresh | Whole | *Sphyrna lewini* | CITES |
| 180 | BW-482 | 3/2/20 | Banyuwangi | BYW7 | PP | Fresh | Whole | *Sphyrna lewini* | CITES |

| No. | ID | Date | Location | Dust Pooled ID Location | Type of Location | Type of Product | Part | Species Identification | CITES Status |
|---|---|---|---|---|---|---|---|---|---|
| 181 | BW-483 | 3/2/20 | Banyuwangi | BYW7 | PP | Fresh | Whole | *Sphyrna lewini* | CITES |
| 182 | BW-484 | 3/2/20 | Banyuwangi | BYW7 | PP | Fresh | Whole | *Sphyrna lewini* | CITES |
| 183 | BW-485 | 3/2/20 | Banyuwangi | BYW7 | PP | Fresh | Whole | *Sphyrna lewini* | CITES |

Notes: Processing plants (PP), export hubs (EH) and an inspector station (AU)

**Table S7.** List of species integrated in the curated reference database and the respective number of individual sequences included per species

| No. | Family Name | Scientific Name | Number of Sequences |
|---|---|---|---|
| 1 | Carcharhinidae | *Carcharhinus amblyrhynchoides* | 5 |
| 2 | Carcharhinidae | *Carcharhinus brevipinna* | 4 |
| 3 | Carcharhinidae | *Carcharhinus falciformis* | 3 |
| 4 | Carcharhinidae | *Carcharhinus leucas* | 2 |
| 5 | Carcharhinidae | *Carcharhinus longimanus* | 2 |
| 6 | Carcharhinidae | *Carcharhinus obscurus* | 2 |
| 7 | Carcharhinidae | *Carcharhinus sorrah* | 3 |
| 8 | Carcharhinidae | *Galeocerdo cuvier* | 2 |
| 9 | Carcharhinidae | *Prionace glauca* | 2 |
| 10 | Carcharhinidae | *Rhizoprionodon oligolinx* | 2 |
| 11 | Carcharhinidae | *Carcharhinus albimarginatus* | 1 |
| 12 | Carcharhinidae | *Carcharhinus obscurus* | 2 |
| 13 | Carcharhinidae | *Triaenodon obesus* | 1 |
| 14 | Alopiidae | *Alopias pelagicus* | 2 |
| 15 | Alopiidae | *Isurus oxyrinchus* | 2 |
| 16 | Alopiidae | *Isurus paucus* | 2 |
| 17 | Alopiidae | *Lamna nasus* | 1 |
| 18 | Sphyrnidae | *Sphyrna lewini* | 5 |
| 19 | Sphyrnidae | *Sphyrna mokarran* | 3 |
| 20 | Sphyrnidae | *Eusphyra blochii* | 1 |
| 21 | Hemigaleidae | *Hemigaleus australiensis* | 2 |
| 22 | Hemigaleidae | *Hemipristis elongata* | 1 |
| 23 | Hemiscylliidae | *Chiloscyllium indicum* | 1 |
| 24 | Hemiscylliidae | *Chiloscyllium punctatum* | 2 |
| 25 | Hemiscylliidae | *Chiloscyllium plagiosum* | 3 |
| 26 | Squalidae | *Squalus hemipinnis* | 1 |
| 27 | Pseudocarchariidae | *Pseudocarcharias kamoharai* | 1 |
| 28 | Stegostomatidae | *Stegostoma fasciatum* | 2 |
| 29 | Triakidae | *Mustelus manazo* | 1 |
| 30 | Dasyatidae | *Himantura gerrardi* | 1 |
| 31 | Dasyatidae | *Neotrygon orientalis* | 2 |
| 32 | Dasyatidae | *Telatrygon zugei* | 2 |

| No. | Family Name | Scientific Name | Number of Sequences |
| --- | --- | --- | --- |
| 33 | Dasyatidae | *Hemitrygon bennettii* | 2 |
| 34 | Dasyatidae | *Himantura leoparda* | 4 |
| 35 | Dasyatidae | *Taeniura lymma* | 1 |
| 36 | Myliobatidae | *Mobula tarapacana* | 1 |
| 37 | Myliobatidae | *Mobula birostris* | 1 |
| 38 | Myliobatidae | *Mobula mobular* | 4 |
| 39 | Rhynchobatidae | *Rhynchobatus australiae* | 2 |
| 40 | Rhynchobatidae | *Rhynchobatus springeri* | 2 |
| 41 | Rhynchobatidae | *Rhynchobatus laevis* | 2 |
| 42 | Pristidae | *Anoxypristis cuspidata* | 2 |
| 43 | Rhinidae | *Rhina ancylostoma* | 2 |
| 44 | Rhinobatidae | *Glaucostegus typus* | 2 |
| 45 | Gymnuridae | *Gymnura poecilura* | 3 |
| | | **Total** | **94** |
